# Fibrin, Bone Marrow Cells and macrophages interactively modulate cardiomyoblast fate

**DOI:** 10.1101/2022.01.06.475189

**Authors:** Inês Borrego, Aurélien Frobert, Guillaume Ajalbert, Jérémy Valentin, Cyrielle Kaltenrieder, Benoît Fellay, Michael Stumpe, Stéphane Cook, Joern Dengjel, Marie-Noelle Giraud

## Abstract

Interactions between macrophages, cardiac cells and the extracellular matrix are crucial for cardiac repair following myocardial infarction (MI). The paracrine effects of cell-based treatments of MI might modulate these interactions and impact cardiac repair. The immunomodulatory capacity of the therapeutic cells is therefore of interest and could be modulated by the use of biomaterials. We first showed that bone marrow cells (BMC) associated with fibrin could treat MI. Then, we interrogated the influence of fibrin, as a biologically active scaffold, on the secretome of BMC and the impact of their association on macrophage fate and cardiomyoblast proliferation.

**Methods:** *In vivo*, two weeks post-MI, rats were treated with epicardial implantation of BMC and fibrin or sham-operated. High-resolution echocardiography was performed to evaluate the heart function and structure changes after 4 weeeks. Histology and immunostaining were performed on harvested hearts. *In vitro*, BMC were first primed with fibrin. Second, non-polarized macrophages were differentiated toward either pro-inflammatory or anti-inflammatory phenotypes and stimulated with the conditioned medium of fibrin-primed BMC (F-BMC). Proteomic, cytokine levels quantification, and RT-PCR were performed. EdU incorporation and real-time cell analysis assessed cell proliferation.

**Results:** The epicardial implantation of fibrin and BMC reduced the loss of cardiac function induced by MI, increased wall thickness and prevented the fibrotic scar expansion. After 4 and 12 weeks, the infarct content of CD68^+^ and CD206^+^ was similar in control and treated animals. In vitro, we showed that fibrin profoundly influenced the gene expression and the secretome of BMC, simultaneously upregulating both pro- and anti-inflammatory mediators. Furthermore, the conditioned medium from F-BMC significantly increased the proliferation of macrophages in a subsets dependent manner and modulated their gene expression and cytokines secretion. For instance, F-BMC significantly downregulated the expression of *Nos2*, *Il6* and *Ccl2/Mcp1* while *Arg1*, *Tgfb* and *IL10* were upregulated. Interestingly, macrophages educated by F-BMC increased cardiomyoblast proliferation.

In conclusion, our study provides evidence that BMC/fibrin-based treatment lowered the infarct extent and improved cardiac function. The macrophage content was unmodified when measured at a chronic stage. Nevertheless, acutely and *in vitro*, the F-BMC secretome promotes an anti-inflammatory response that stimulates cardiac cell growth. Finally, our study emphases the acute impact of F-BMC educated macrophages on cardiac cell fate.

## Introduction

Macrophages, the hallmark for tissue healing and wound formation, mediate the inflammatory response following myocardial infarction (MI) and contribute to its resolution (1). Frangogiannis et al. (2) primarily showed that shifting the balance of macrophages from an inflammatory to an anti-inflammatory phenotype is essential for cardiac regeneration. The roles of macrophages are multifaced, and their importance in the regeneration of the neonatal heart has been identified (3), while it remains mostly unrevealed in the adult myocardium. Extremely versatile, macrophages adopt a variety of functional phenotypes depending on signals in their environment ranging from pro-inflammatory to anti-inflammatory and pro-resolution phenotypes (4). The different subsets of macrophages can be defined by their function, expression of genes, or responsiveness to activating cytokines (5). Two major types, namely, alternatively- and classically-activated macrophages, are obtained by treatment of unpolarised macrophages, respectively, with Interleukin 4 (IL4) or lipopolysaccharide (LPS) associated with interferon (INF) respectively.

The spatiotemporal interaction of macrophages with the cardiac extracellular matrix (ECM), cardiomyocytes and non-cardiomyocytes are believed to be a key feature in cardiac repair and regeneration (6–8). Consequently, manipulating macrophage subsets has become an emerging therapeutic strategy in numerous diseases, including cardiovascular diseases (CVD) (9, 10). Due to their immunomodulatory capacities, cell-based therapies, including bone-marrow cells (BMC), mononuclear cells (MNC) (11), mesenchymal stromal cells (MSC)(12, 13), anti-inflammatory macrophages (14) or their secreted extracellular vesicles have gained interest for treating CVD such as MI (12, 15, 16). These cell secretome, composed of cytokines, growth factors, and extracellular vesicles or exosomes, is accountable for their paracrine effect that improves cardiac structural and functional outcomes accompanied by mobilization and polarisation of macrophages (11, 17, 18). For instance, Vagnozzi et al. (11) demonstrated that seven days after ischemia/reperfusion (I/R), bone marrow MNC and MSC improved the heart function through an acute immune response characterized by the mobilization of CCR2^+^ or CX3CR1^+^ macrophages. In addition, Deng al. al (17) promoted anti-inflammatory macrophage polarisation with MSC-exosomes and demonstrated amelioration of cardiac damage after MI.

The use of biomaterials combined with cells is recommended for cell-based therapies (19). The scaffolds foster cell retention and survival. Nevertheless, scaffolds may also modulate macrophage phenotypes depending on their composition and structure. (20, 21). Fibrin, a biologically active scaffold, has gained an increasing interest in tissue engineering (22). Its general use interrogates its influence on the fate of cells. In the present study, fibrin combined with unfractionated BMC was administered sub-chronically in a rat model of MI. Beneficial functional outcomes were recorded after four weeks and were associated with a reduced fibrotic scar. In vitro investigations revealed that the secretome of fibrin primed-BMC (F-BMC) specifically activated macrophages with anti-inflammatory and mitogenic properties.

## Results

### In vivo study

#### BMC associated with fibrin restores the cardiac function loss and reduces the fibrotic scar in the infarct heart

Epicardial implantation of fibrin and BMC was performed two weeks after the induction of MI by left anterior descending coronary ligation in a rat model, and the therapeutic potential was evaluated.

Before the treatment and four weeks post-treatment, functional and structural characteristics were assessed by high-resolution echocardiography. The differences in ejection fraction (EF) and fractional shortening (FS) in the treated group were calculated. As shown in figure 1A, the treated group showed statistically significant EF and FS gains, compared to the untreated group that showed a loss in cardiac functions (figure 1A). It is important to note that the gain in EF and FS showed high inter-individual variability. Indeed, the highest and the lowest EF gain measured were +14% and -8%, respectively, in the treated groups and +3% and -11% in the control group. Structural adaptations also differed between groups with augmented systolic wall thicknesses and reduced LV volumes in treated animals (figure 1A).

**Figure 1A:**
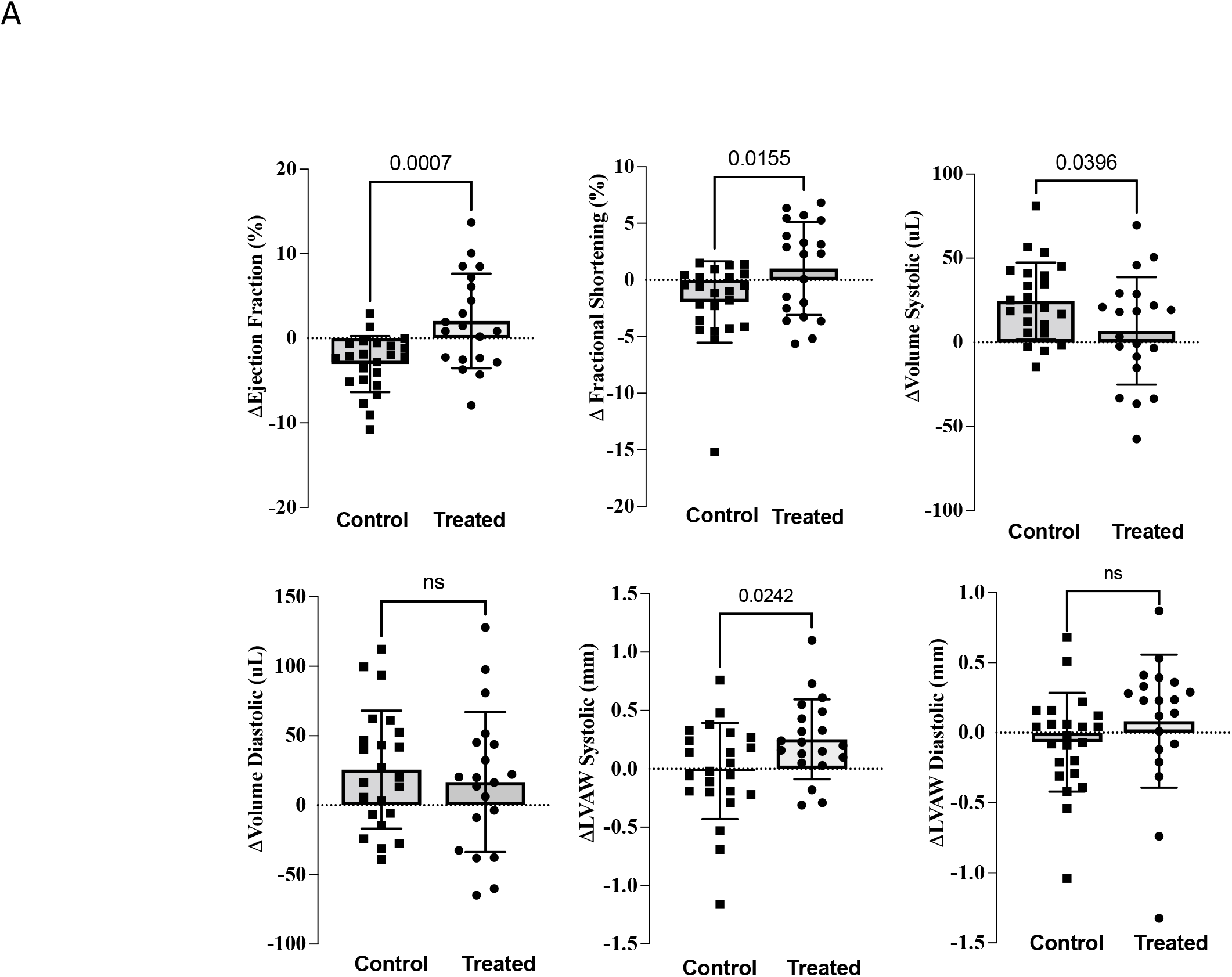
Fibrin and BMC treatment reduces systolic heart function loss. Cardiac function was assessed by high-resolution echocardiography. The changes (Δ) in Ejection Fraction, Fractional shortening, left ventricle (LV) volumes and LV wall thickness (LVAW) were calculated as the difference between 4 weeks post- and pre-treatment. The control group is untreated (sham) infarcted animals (n=23); the treated group is animals treated with epicardial fibrin and BMC (n=20).

**Figure 1B:**
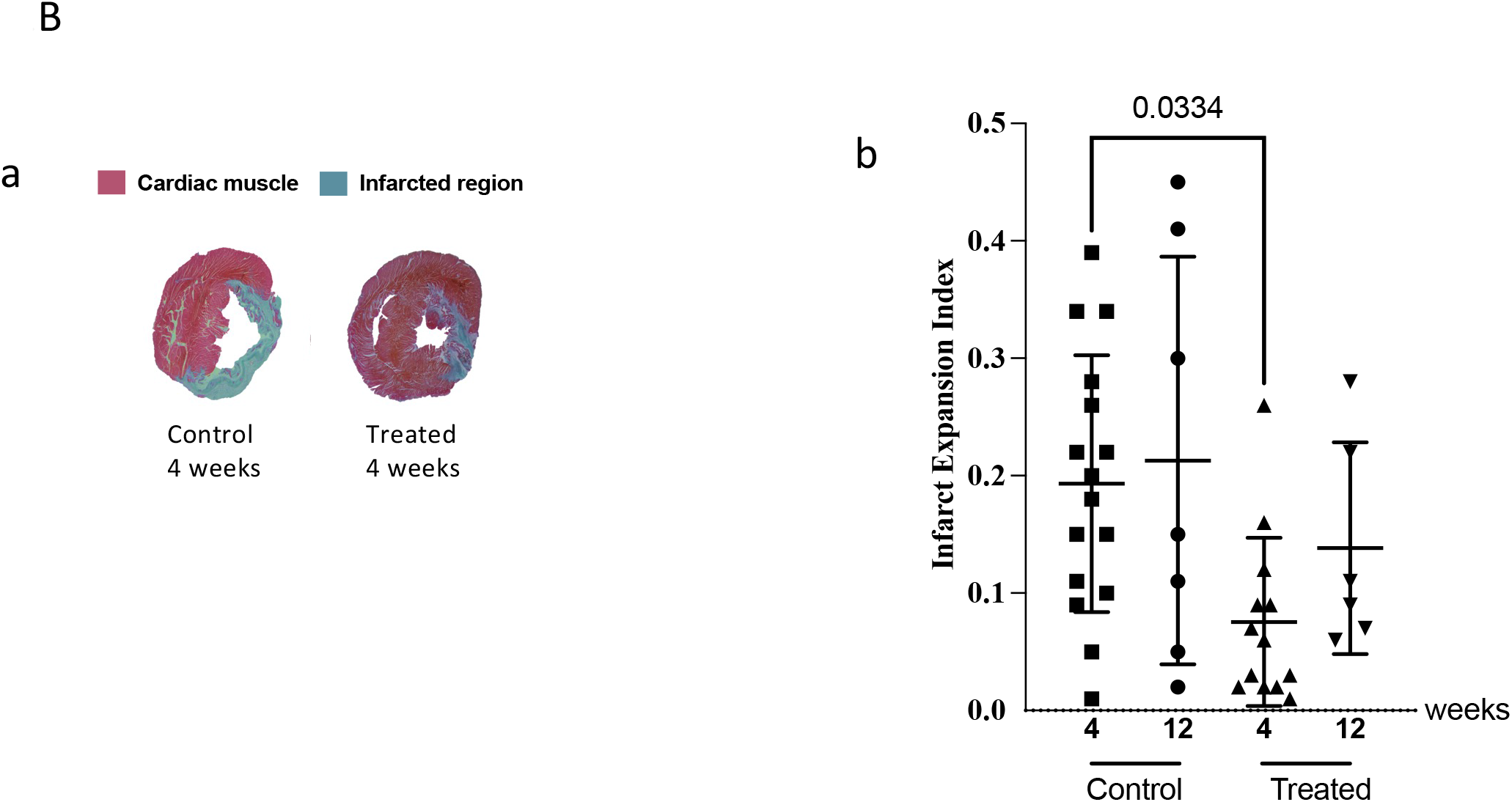
Fibrin and BMC treatment reduces fibrotic scar. (a) Representative Masson-Goldner trichrome stained heart cross-sections from treated and control (sham) animals, 4 weeks post-treatment. The fibrotic scar tissue is in blue; in red is the remote tissue. (b) Infarct Expansion Index was measured on histologic sections 4 weeks or 12 weeks post-treatment. 5 to 6 cross-sections from a systematic sampling of the whole heart were averaged for each animal. Each point represents one animal. The Control group is infarcted animals that received sham treatment. The treated group is animals treated with epicardial fibrin and BMC. The values are shown mean ± SD

Furthermore, Masson’s trichrome staining was performed four weeks post-treatment (figure 1B). calculation of the infarct expansion index demonstrated that the scar was substantially smaller in treated than in the untreated groups (figure 1B), indicating that the treatment significantly decreased cardiac fibrosis. Seven animals per group were kept twelve weeks post-treatment. One animal in the treated group died after 11 weeks. Twelve weeks post-treatment, the mean scar expansion index was decreased in treated animals compared to control ones; however, the difference did not reach statistical significance.

Further immunostaining showed that the percentage of CD68 and CD206 macrophages present in the infarcted myocardium and the peri infarcted area after 4 and 12 weeks was similar in all groups (Figure S1).

### In vitro study

#### Unique characteristics of F-BMC, including growth, gene expression and secretion profile, distinguish them from BMC

BMC are a heterogeneous population of cells as indicated by markers from the mesenchymal and hematopoietic lineages (Figure S2); the mesenchymal CD90^+^ cells were most abundant. Fibrin induced a change in the morphology of BMC (Figure 2). While BMC displayed a homogeneous spindle-like shape, F-BMC presented rounded cell morphology. The proliferation of F-BMC was measured by EdU incorporation and revealed a significant reduction compared to BMC (Figure 3A). In addition, cell growth was assessed using a real-time cell analyzer (RTCA), in which microelectrodes measure the electrical impedance of the cell populations assessing both, the cell number and cell spreading. Results are presented as the area under the curve (AUC) after 5 days (Figure 3B, respectively). The AUC of F-BMC was significantly lower than the AUC of BMC, confirming a reduced cell growth and spreading. Taking all together, our data show that fibrin decreased both the proliferation and the spreading of BMC.

**Figure 2:**
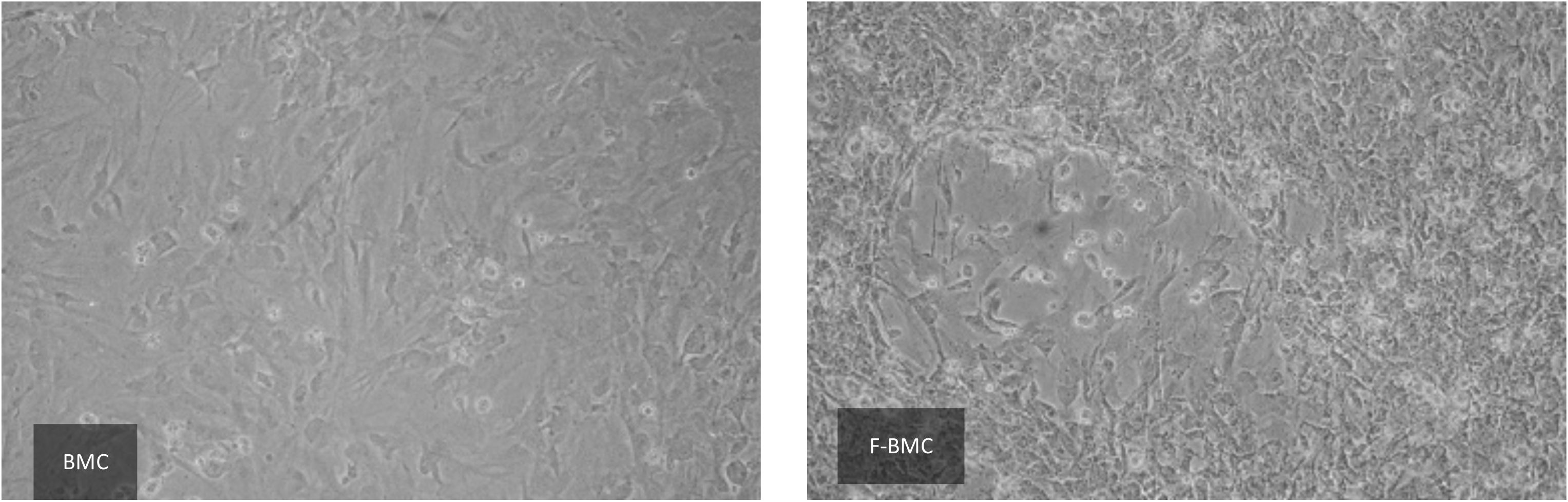
Fibrin modulates the BMC morphology: Representative pictures of BMC cultured with fibrin (F-BMC) or without (BMC) showing a heterogeneous population of cells with different morphology and spreading.

**Figure 3:**
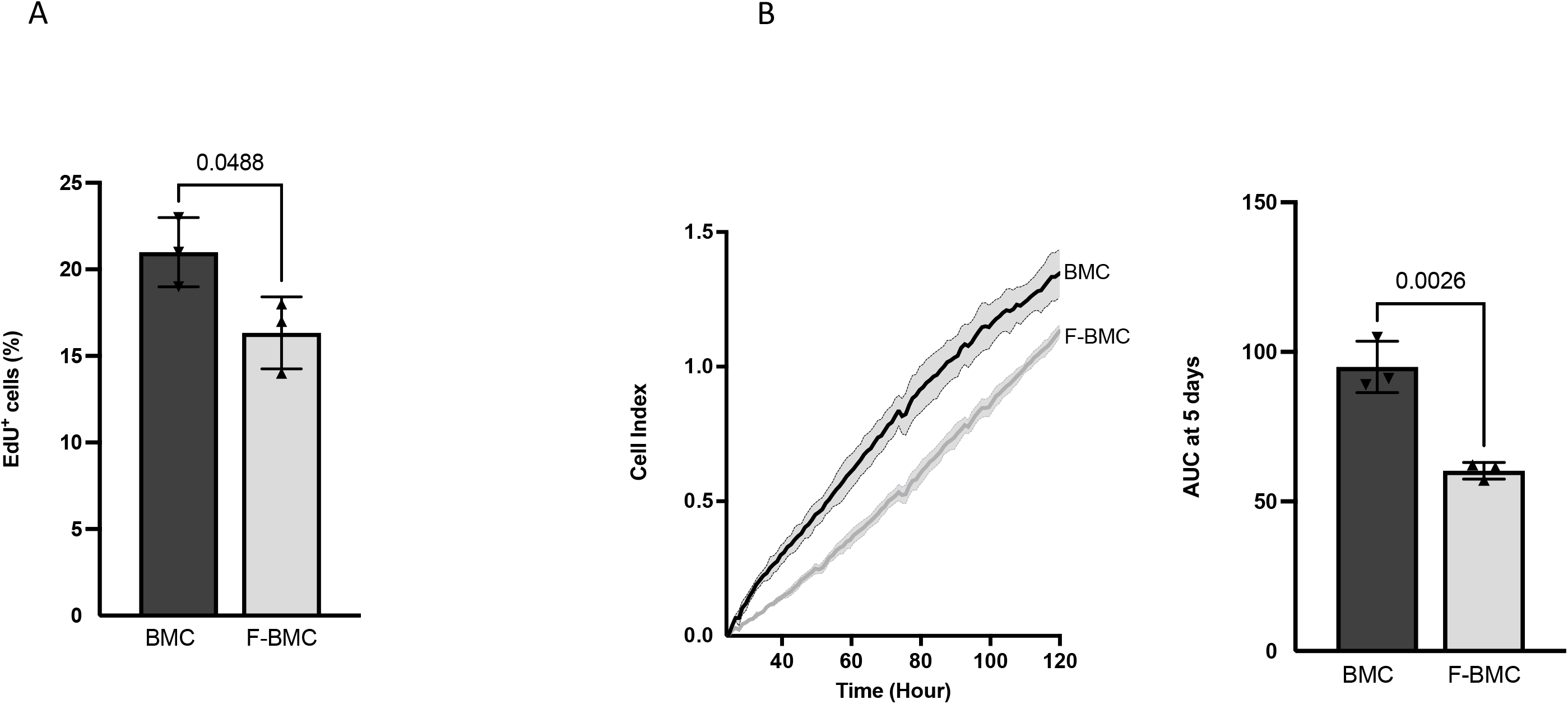
Fibrin modulates BMC proliferation: BMC were cultured with (F-BMC) or without fibrin (BMC). Cell growth was assessed by (A) EdU incorporation during 48h and by (B) RTCA: Cell index was measured over 120 hours, plotted, and the AUC calculated. The values are shown mean ± SD; n=3 biologically independent samples of pools the BMC of 3 rats.

The Conditioned media (Cd-media) of F-BMC and BMC were analyzed by liquid chromatography (LC)-tandem mass spectrometry (MS/MS). In total, 1712 proteins were identified (protein and peptide FDR<0.01), of which 769 were quantified in minimally two replicates out of three per group using label-free quantification based on respective peptide ion currents (23) . Hierarchical clustering of data nicely discriminated the two experimental groups indicating that Cd-media of F-BMC and BMC differed in their proteomic composition on a global scale (Figure 4A). In total, protein abundances of 339 proteins were significantly altered between the two groups, 185 were significantly upregulated, and 154 were downregulated comparing F-BMC to BMC (Figure 4B; t-test, FDR<0.05). Upregulated proteins could be linked to metabolic and catabolic processes, inflammation response and ECM remodelling (Figure 4C, Supplemental Table 1; BH FDR<0.02).

**Figure 4:**
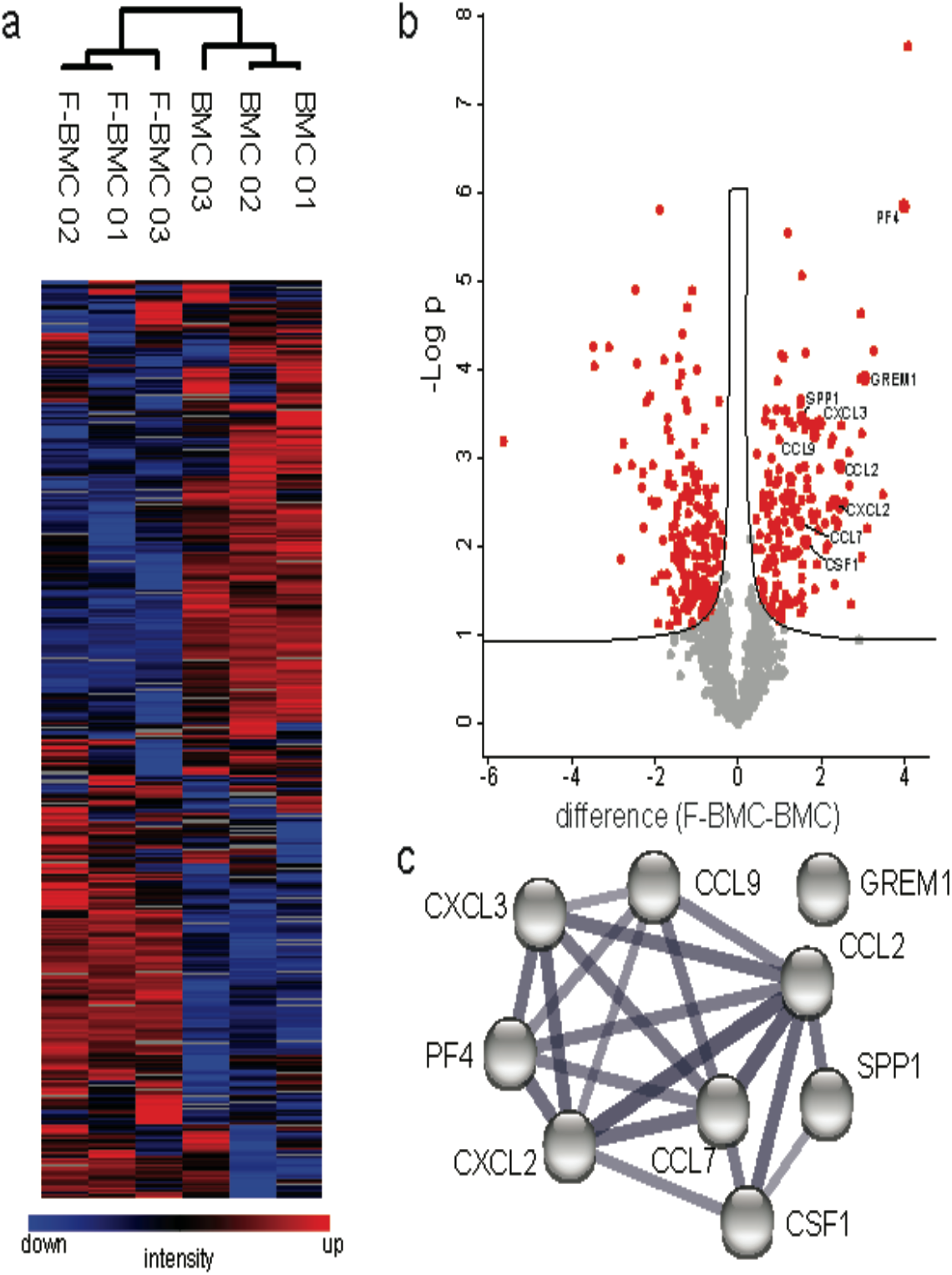
Expression proteomic analyses of BMCs cultured with and without fibrin-based substrate. (a) Hierarchical clustering of protein abundances using log2 transformed and z-normalized LFQ intensities indicates global alterations of CM due to culture conditions. Grey squares indicate proteins not detected in respective samples. (b) Volcano plot analysis highlights significantly altered proteins due to culture condition in red (FDR<0.05). The black line indicates S0 of 0.1. (c) Eight cytokines known to interact based on STRING DB are significantly downregulated in BMCs cultured with the fibrin-based substrate (58). Thickness of edges indicates confidence of data support.

In complement, RT-PCR analyses of related genes were performed on cell lysates. Differential expressions of genes involved in inflammation and ECM organization were prioritized (Figure 5). Furthermore, as proteomics did not allow quantifying interleukins secretion, their expressions were investigated by RT-PCR. Fibrin priming induced a significant upregulation of *TIMP1* expression and down-regulation of the proteolytic enzymes *MMP3* and *MMP9 (*Figure 5). In agreement, the level of protein secretions of TIMP1 and MMP9 exhibited the same significant trends (Supplemental Table 1). *Vcam1*, *Icam* and the inflammation mediators *Nos2* and *Tgfb* were significantly downregulated by fibrin, whereas Il*1ra*, *Il1b*, *IL6,* and *Gmcsf* expression increased. (Figure 5). Concerning *Tnfa,* the gene expression was upregulated; however, the difference in the secretion of TNF-alpha between BMC and F-BMC did not reach statistical significance as shown in the proteomic analysis (Supplemental Table 1). *Ccl2/Mcp1* gene expression and related protein secretion were augmented (Figure 5, Supplemental Table 1). While *Csf1/Mcsf* expression was not statistically changed, the secretion of the protein increased. Chemokines were also significantly modulated by fibrin. *Ccl5/Rantes* expression increased while *Cxcl10* decreased (Figure 5). In parallel, the secretion of CxCL2, CxCL3, CxCL4/PF4, CCL7, CCL9 were significantly higher in F-BDMC (Supplemental Table 1). Also, proteomic analysis revealed the modulation of proteins such as Osteopontin (ONP/SPP1) and Gremlim1 (GREM1) that were both increased by fibrin (Supplemental Table 1).

**Figure 5:**
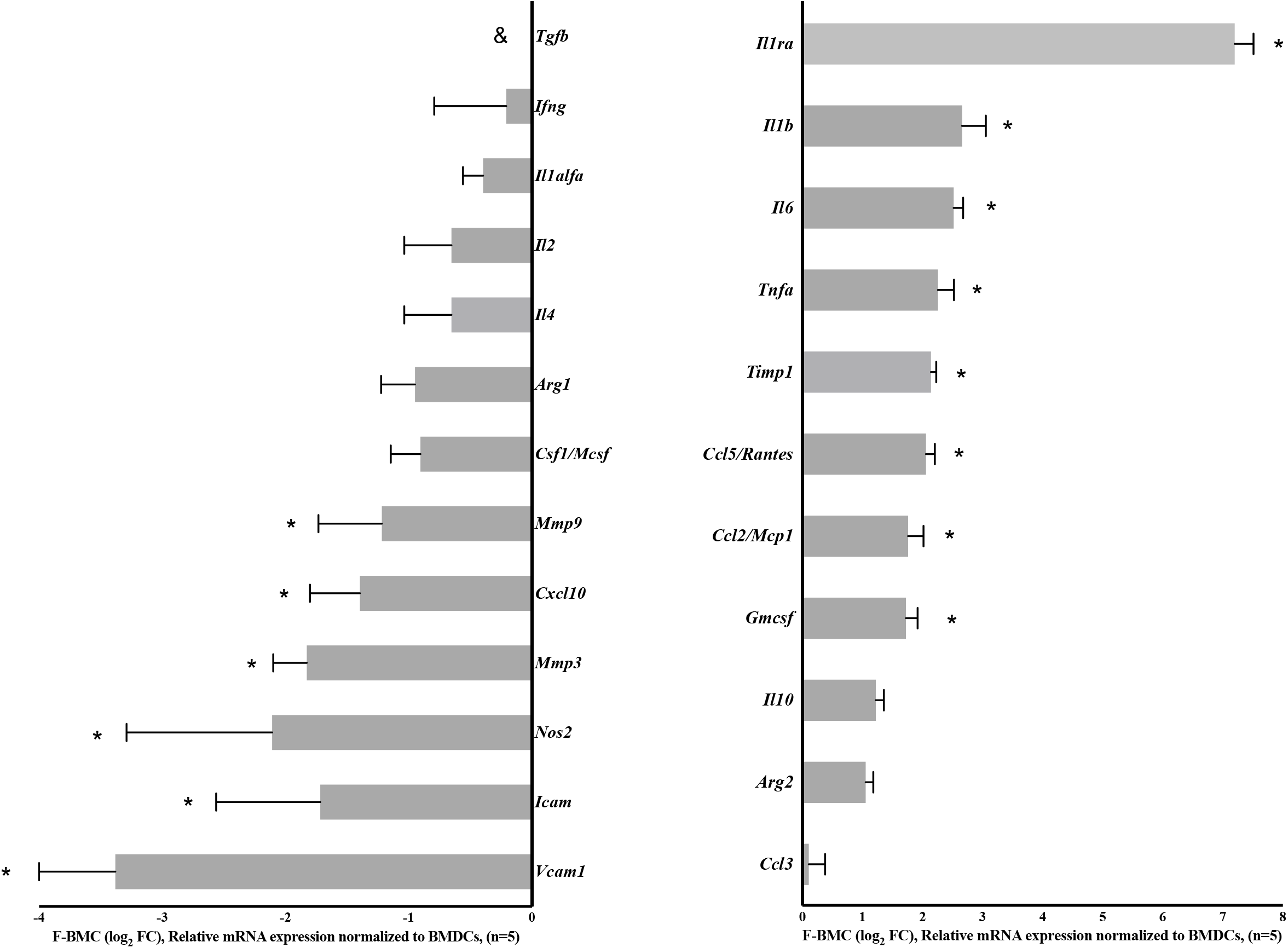
Fibrin modulates the BMC gene expression. Real-time PCR measurement of selected genes presented as fold change (FC). n=5 biologically independent samples of cell pools from 3 rats. *p < 0.05, shows the statistical significance between differential gene expression of F-BMC related to BMC assessed by one-way ANOVA and Dunnett’s test.

Collectively, these results show that fibrin profoundly influenced gene expression and the cell proteome, simultaneously upregulating both pro- and anti-inflammatory mediators.

### F-BMC secretome promotes the proliferation of undifferentiated and anti-inflammatory macrophages

The effects of the secretomes of BMC and F-BMC on macrophage fates were investigated. The proliferation of undifferentiated (M_(-)_), pro-and anti-inflammatory macrophages (respectively, M_(LPS, IFN)_ and M_(IL4)_) cultured with Cd-media from BMC, F-BMC and fibrin were assayed by EdU incorporation. After 48 hours, the percentage of EdU^+^ M_(-)_ increased significantly when cultured with Cd-media from both F-BMC and BMC (Figure 6A) relative to the unconditioned medium (control). Cd-medium from fibrin had no significant effect. The proliferation rate of M_(LPS, IFN)_ remained quantitatively low and similar for all conditions (Figure 6B). For M_(IL4),_ Cd-medium from F-BMC significantly increased their proliferation rate (Figure 6C) compared to unconditioned medium, while Cd-media from BMC and fibrin did not significantly differ. In addition, the proliferation of M_(IL4)_ was statistically significantly more stimulated by F-BMC secretome than by the BMC one. Altogether, our results show that fibrin-primed BMC modulated macrophages proliferation; in particular, the mitogenic properties of the Cd-media were utmost for F-BMC educated M_(IL4)_.

**Figure 6.**
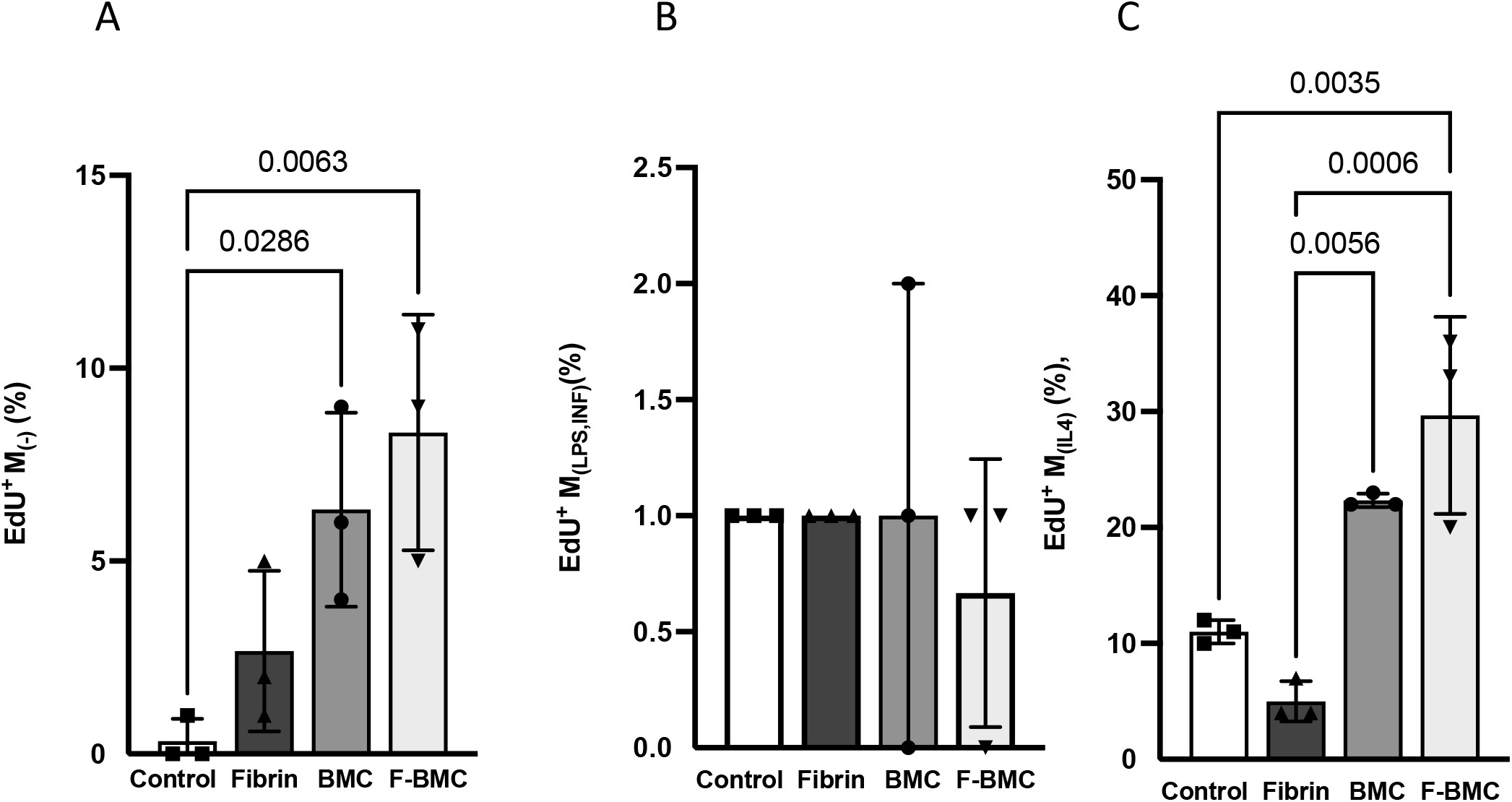
F-BMC and BMC conditioned media modulate macrophage proliferation. The proliferation of the different macrophages phenotypes was assessed by EdU incorporation. (A) M_(-),_ (B) M_(LPS,IFN),_ and (C) M_(IL4)_ macrophages were cultured with unconditioned medium (Control) or conditioned media from with Fibrin, F-BMC and BMC. The values shown are mean ± SD. n= 3 biologically independent pools of macrophages from 3 animals, therefore, in total, 9 animals per group.

### F-BMC secretome induces a macrophage phenotype switch

RT-PCR analyses were performed on lysates of macrophages cultured with Cd-media from F-BMC (F-BMC educated macrophages) (Figure 7), BMC and fibrin (Figure S3). F-BMC-educated M_(-)_ showed a significantly downregulated expression of pro-inflammatory genes, particularly *Nos2*, *Il6* and *Ccl2/Mcp1*. Anti-inflammatory genes such as *Arg1, Tgfb* and *IL10* were significantly upregulated (Figure 7A). Similar gene regulations were recorded for F-BMC-educated M_(LPS,INF)_ with a significant downregulation of Il1b and no change for *Ccl2/Mcp1* (Figure 7B). Remarkably, the anti-inflammatory phenotype of the macrophage M_(IL4)_ was further stimulated. All tested pro-inflammatory genes were significantly downregulated, and all studied anti-inflammatory ones were upregulated (Figure 7C).

**Figure 7.**
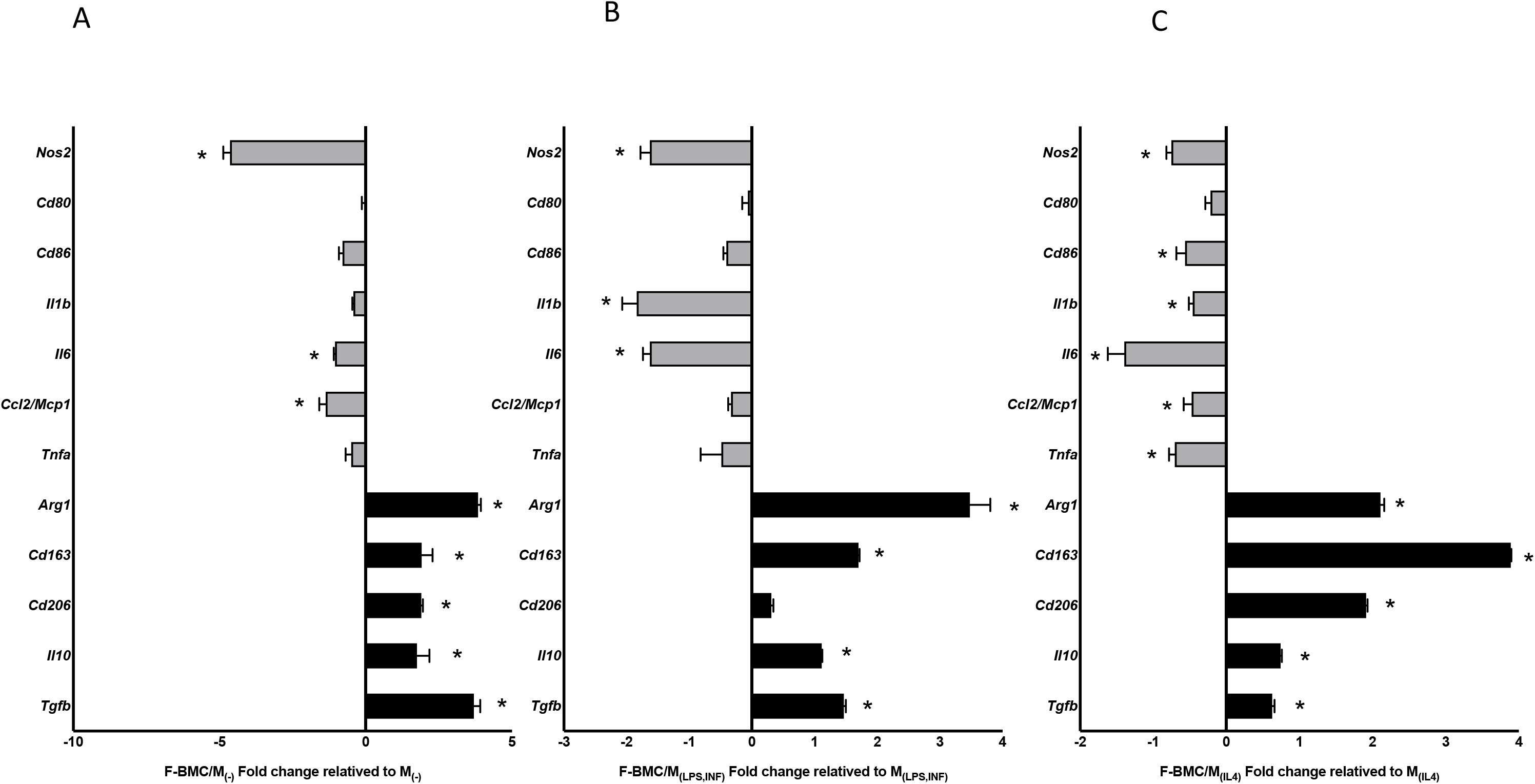
F-BMC conditioned medium modulates the macrophage expression profile. Relative gene expression of pro-inflammatory markers (grey) and anti-inflammatory markers (black) measured in F-BMC educated macrophages: (A) M_(-)_, (B) M_(LPS, IFN)_ and (C) M_(Il4)_ relative to uneducated ones. The values shown are mean ± SD. All n= 3 biologically independent samples constituted of macrophages pools, each pool was obtained from 3 animals, therefore, in total, 9 animals per group. *p < 0.05, shows the statistical significance between differential gene expression of F-BMC educated macrophages and uneducated ones assessed by one-way ANOVA and Dunnett’s test.

The expressions of the surface markers of F-BMC-educated macrophage were significantly altered, specifically the markers *Cd206* and *Cd163* related to alternatively activated macrophage phenotypes. *Cd206* was significantly upregulated in M_(-)_ and M_(IL4)_ (Figure 7A and 7C), while *Cd163* was significantly upregulated in all macrophages. Further, classical activated macrophage marker *Cd86* was downregulated in M_(IL4)_. (Figure 7B). The switch of macrophage plasticity to an anti-inflammatory profile was observed in BMC educated macrophages. The gene expression switch was more prominent for F-BMC educated M_(-)_ and M_(LPS, IFN),_ (Figure S3). Fibrin also showed a different regulatory pattern in M_(LPS,INF)_ and M_(-)_ compared to BMC and F-BMC (Figure S3).

The levels of IL-1beta, IL-6 and TNF-alpha secreted by F-BMC-educated macrophages were further quantified by Enzyme-linked immunosorbent assay (ELISA). Unconditioned medium served as control. IL-1beta levels were significantly decreased in M_(LPS,INF)_ and M_(IL4)_ with respective -2.8 and -5.8 fold changes compared to control (Figure 8). IL-6 levels were significantly decreased for M_(-)_, M_(LPS,INF)_ and M_(IL4)_ with -3.7, -2.8 and -3.0 fold changes, respectively (Figure 8). TNF-alpha secretion was also significantly lower in M_(-)_ and M_(LPS,IFN)_ with respective fold changes of -4.0 and -2.8 (Figure 8). The decrease in the secretion of cytokines corroborated the changes in gene expression measured with RT-PCR. Taking all together, the results show that F-BMC-educated macrophages decreased both the expression and the secretion of different proteins related to a pro-inflammatory phenotype.

**Figure 8.**
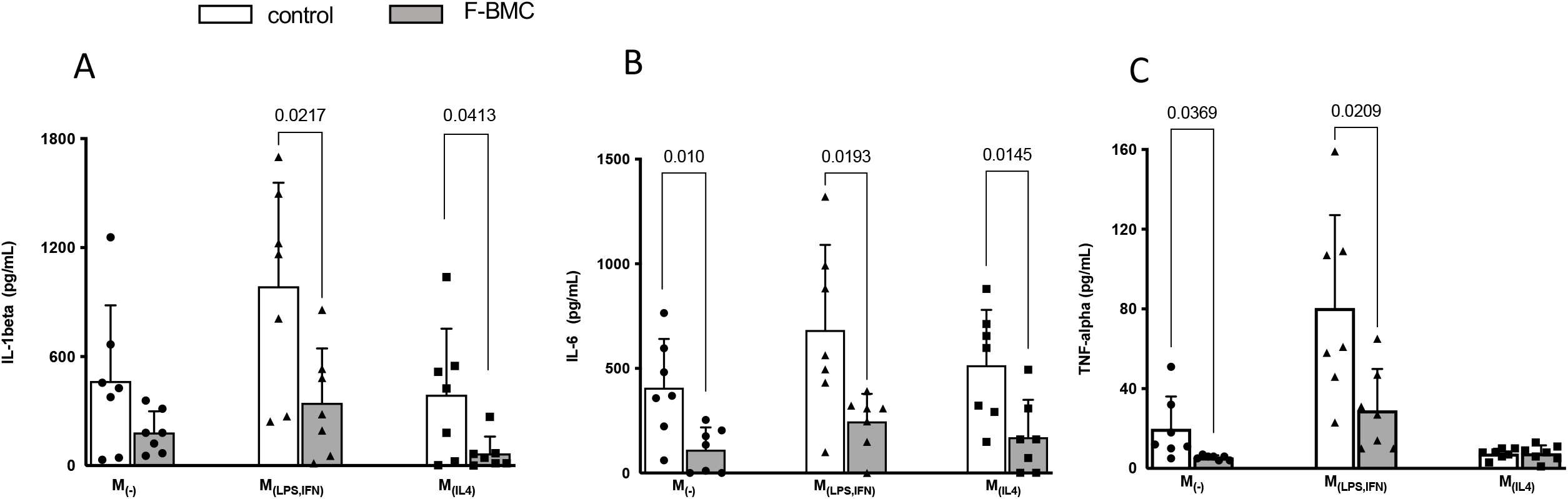
F-BMC conditioned medium modulated the macrophage secretion profile. Cytokine expression levels of (A) IL-1 beta, (B) IL-6, and (C) TNF-α were quantified by ELISA in cell culture supernatants of M_(-)_, M_(LPS,IFN),_ and M_(IL4)_ educated with F-BMC conditioned medium or unconditioned medium. The values shown are mean ± SD. n= 7 biologically independent macrophages pools; each pool was obtained from 3 animals.

### F-BMC secretome promotes cardiomyoblast spreading

Cardiomyoblasts (H9C2) was chosen as a model of proliferative cardiac cells. The proliferation of H9C2 cultured with Cd-media from BMC, F-BMC and fibrin was compared to an unconditioned medium (control) and measured using EdU incorporation. After two days, the percentage of EdU^+^ H9C2 was significantly higher with F-BMC compared to BMC and Fibrin Cd-media (Figure 9A) and similar to the control.

**Figure 9.**
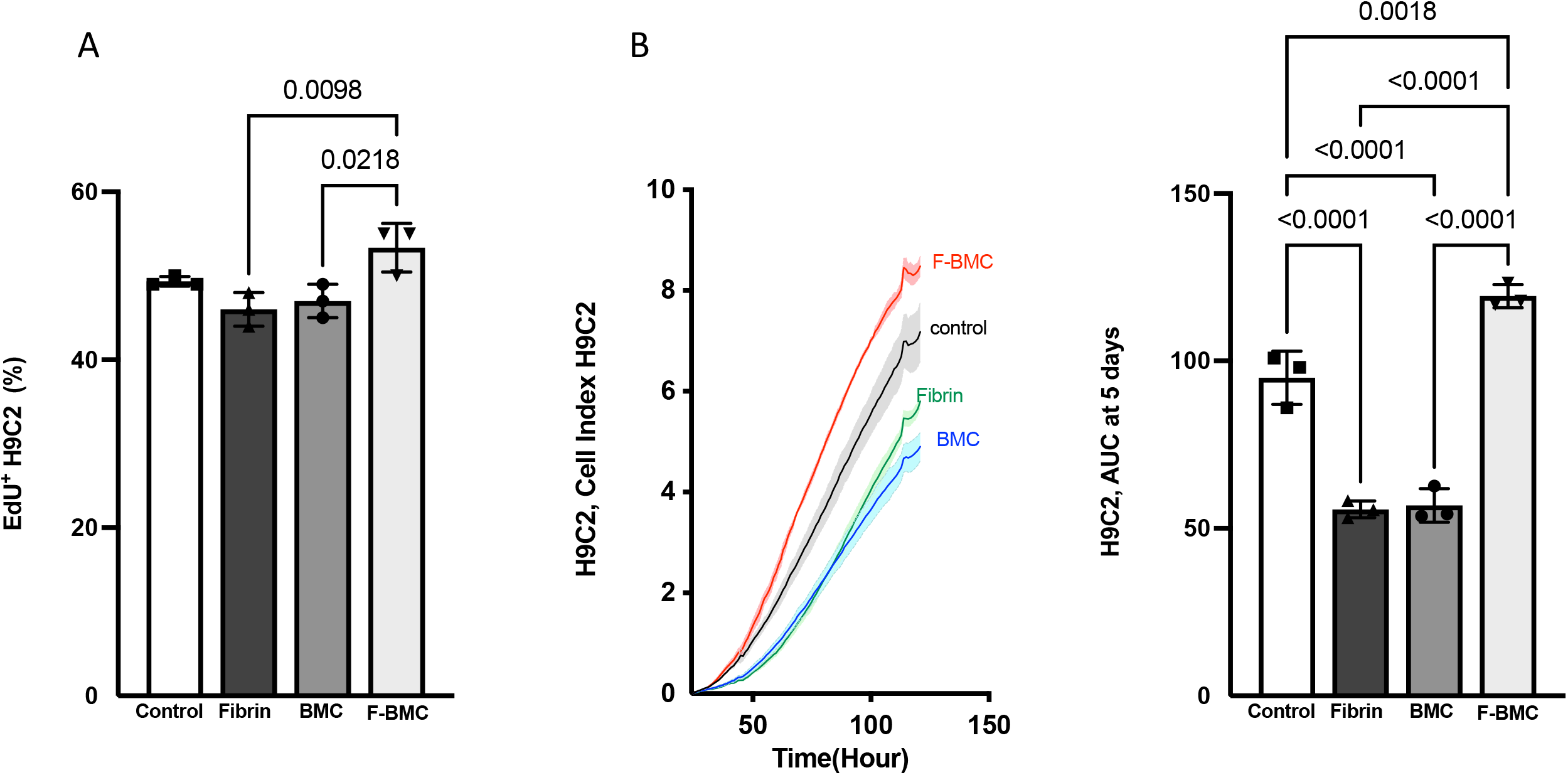
F-BMC, BMC and fibrin secretomes altered cardiomyoblasts H9C2 growth. H9C2 were cultured with Cd-media from F-BMC, BMC or fibrin. Standard growth medium served as control. H9C2 growth was assessed by (A) EdU incorporation during 48h and by (B) RTCA: Cell index was measured over 120 hours, and the AUC was calculated after five days. The values are shown mean ± SD; n=3 biologically independent experiments.

In addition, the F-BMC secretome significantly increased the AUC of H9C2, suggesting an increased spreading with or without an increased cell number. The cell index measured by RTCA (Figure 9B) corroborated the proliferation assay. Taking all together, our results suggest that F-BMC secretome induced an increase in H9C2 spreading and had no effect on cell proliferation when compared to the control condition. On the opposite, fibrin and BMC secretomes reduced both H9C2 spreading and proliferation compared to F-BMC.

### Alternatively-activated macrophages promote cardiomyoblast proliferation

Cd-media obtained from unpolarised and polarised macrophages were used to culture H9C2 (Figure 10). EdU^+^ cells and AUC were significantly increased for M_(IL4),_ suggesting that the secretome of alternatively activated macrophages increased the H9C2 proliferation rate. M_(-)_ and M_(LPS,INF)_ did not alter the spreading of H9C2 compared to control. M_(LPS,INF)_ reduced H9C2 proliferation.

**Figure 10.**
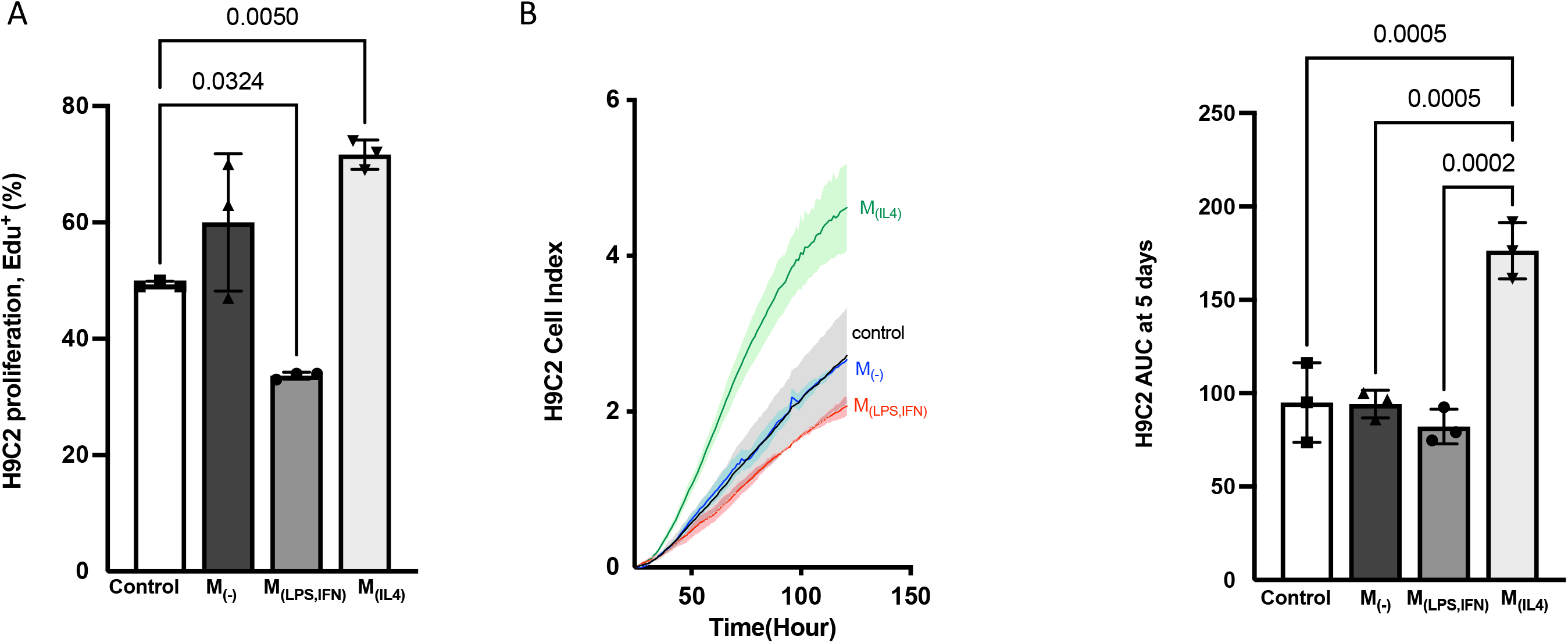
Macrophage secretomes altered cardiomyoblasts H9C2 proliferation rate and spreading. H9C2 were cultured with Cd-media from M_(-)_, M_(LPS,IFN)_ or M_(IL4)_. Standard growth medium served as control. H9C2 growth was assessed by (A) EdU incorporation during 48h and by (B) RTCA: Cell index was measured over 120 hours, and the AUC was calculated after five days. The values are shown mean ± SD; n=3 biologically independent experiments.

### F-BMC-educated-macrophages demonstrate paracrine mitogenic properties on cardiac cells

Educated macrophages were obtained from M_(-)_, M_(LPS,IFN)_ or M_(IL4)_ macrophages primed with Cd-media from BMC, F-BMC or fibrin. Then, H9C2 were cultured with secretomes from educated or uneducated macrophages (Figure 11). The secretome of F-BMC-educated M_(-)_ (labelled as M_(-)_/F-BMC) modulated the growth of H9C2 as shown by the significant increase of the percentage of EdU^+^ H9C2 (Figure 11A) and the AUC at 5 days (Figure 11B) compared to the uneducated M_(-)_ secretome.

**Figure 11.**
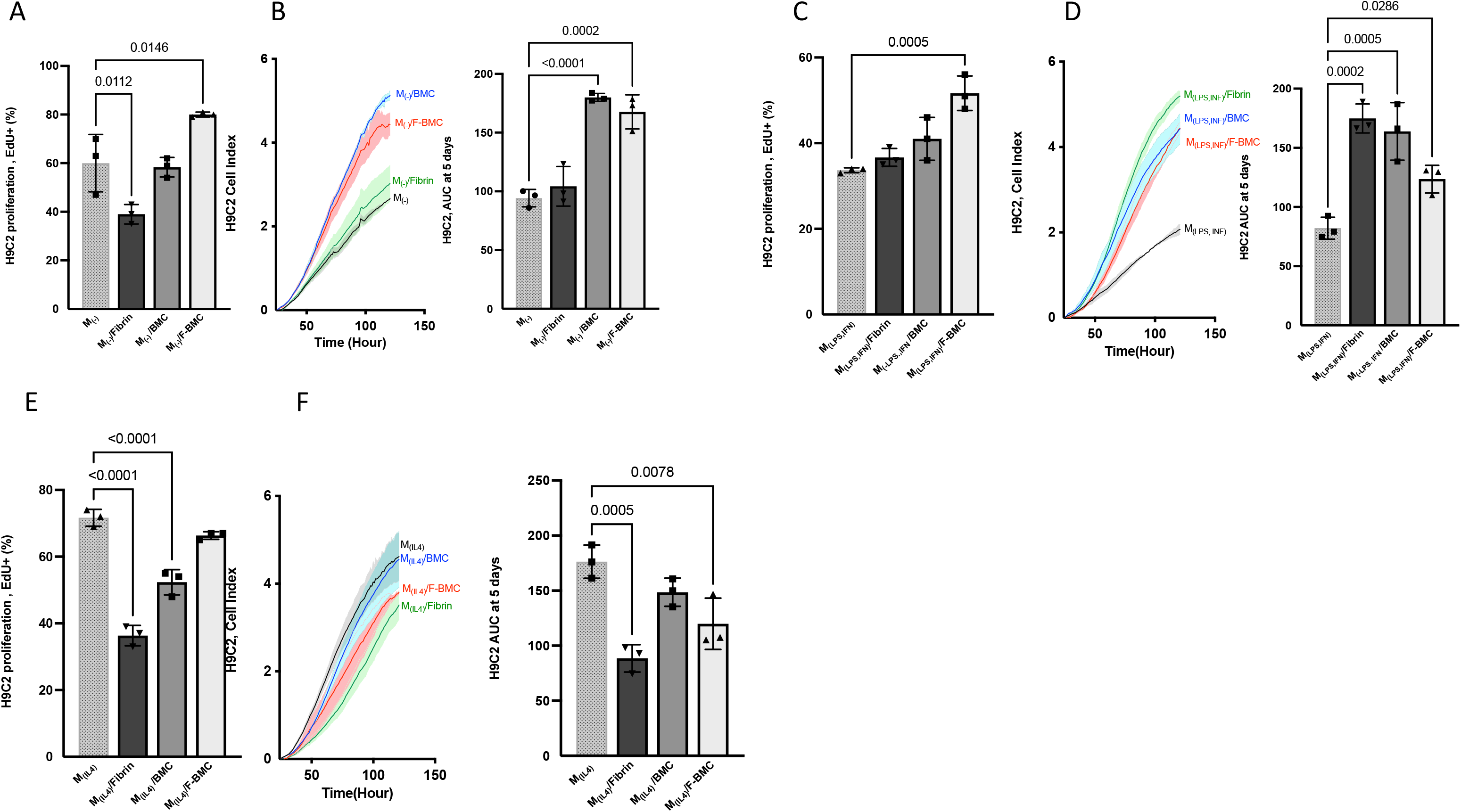
The conditioned media of educated macrophages altered cardiomyoblasts H9C2 proliferation rate and spreading. H9C2 were cultured with Cd-media from (A and B) educated M_(-)_, (C and D) educated M_(LPS,IFN)_ or (E and F) educated M_(IL4)_.macrophages were cultured with respectively Cd -media from fibrin, BMC or F-BMC. Cd medium from uneducated served as control. H9C2 growth was assessed by (A, C and E) EdU incorporation during 48h and by (B, D and F) RTCA: Cell index was measured over 120 hours, and the AUC was calculated after five days. The values are shown mean ± SD; n=3 biologically independent experiments.

Similar trends were observed for M_(LPS,INF)_ /F-BMC (Figure 11C and 11D) while M_(IL4),_ /F-BMC showed different effects. Indeed, M_(IL4)_/F-BMC induced proliferation of H9C2 similar to the control (uneducated M_(IL4)_) and a great reduction in H9C2 spreading (Figure 11E and 11F). Notably, the uneducated M_(IL4)_ induced a strong proliferation of H9C2 (Figure 11E and 11F). This proliferation capacity is maintained when M_(IL4)_ are educated with F-BMC while the H9C2 spreading is reduced.

## Discussion

In the present study, fibrin and BMC were combined to treat sub-chronically the ischemic myocardium. Epicardial implantation reduced the MI related loss of cardiac function and prevented the fibrotic scar expansion. The therapeutic effect of cell-based therapy is associated with an acute paracrine activation of in situ repair mechanisms (24). Indeed, secreted mediators are believed to be essential for overcoming post-natal cardiomyocytes cell cycle arrest, manipulating the microenvironment, stimulating progenitor differentiation and promoting the functional polarisation of the non-myocyte cell population.

In addition, following cell-based treatment, the role of the inflammatory response and its resolution has gained increasing interest. The presence of macrophages and their role have recently been explored. Vagnozzi et al. (11) demonstrated that CD68^+^ macrophages were increased within the area of cell injection after three- and seven days post-injection; however, their presence vanished after two weeks (11). The authors suggested that macrophages were associated with the presence of injected cells (11). Nowadays, it is validated that the therapeutic cells survived only a few days after implantation (11, 25). In the present study, MI content of CD68^+^ and CD206^+^ macrophages were similar in treated and untreated groups after 4 and 12 weeks. The presence of therapeutic cells is unlikely, and accordingly, the macrophage content is unlikely to be affected at these late stages. Nevertheless, when MSCs were administrated together with a slow degrading polycaprolactone matrix, our previous study identified the presence of CD68^+^ cells four weeks post-treatment (25), suggesting that the chronic presence of macrophage might also be dependent on the lasting presence of the matrix. In the current study, fibrin, as a rapidly degrading matrix, was absent after 4 weeks, which may explain the lack of difference in the macrophage content.

It is reasonable to suggest that the initiation of cardioprotective events leading to scar reduction would occur rapidly after the treatment. Therefore, to understand these potential early events, we investigated *in vitro* the interaction between the matrix, the therapeutic cells, the macrophages and cells of cardiac origin.

The association of cell and matrix have gained increasing interest as a therapeutic product for cardiac repair (26). The matrix plays a critical role in prolonging cell survival after transplantation (27, 28). Initially used as a scaffold to provide physical substrate for cell survival, the matrix also has multiple effects (29). The present *in vitro* study provides evidence of the impacts of the matrix on the therapeutic cells. The fibrin matrix alters the morphology of BMC, their gene expression and protein secretion and fosters their immunomodulatory capacities.

### Impact of fibrin on BMC and their properties

First, we report an effect on BMC morphology and proliferation. BMC is a heterogeneous population of cells with a prevalence of mesenchymal CD90^+^ cells. It is well established that unfractionated BMCs are composed of monocytes, lymphocytes, hematopoietic stem cells, MSC and progenitors. Accordingly, hybrid properties and inter-cellular cross-talks between all cell types can be expected (30). We demonstrated that fibrin modified the BMC shape, reduced their spreading and proliferation. In agreement, previous studies demonstrated that MSC showed a modified morphology, a reduced size and a decreased proliferation when cultured with fibrin (31, 32). Furthermore, the fibrin-induced BMC morphological changes were associated with a downregulation of *Icam* and *Vcam1*. These adhesion molecules mediate the interaction between cells and the ECM, and their downregulation could explain a reduced spreading of the cells resulting in a reduced size and rounded shape.

Second, we demonstrated that fibrin induced changes in BMC gene expressions. The most prominent gene upregulation was the expression of *Il1 receptor antagonist* (*IL1ra*) and, to a lesser extent of *Il1b*, while *IL1a* remained unchanged. In normal homeostasis, IL1ra counterbalances the effect of IL1b and IL1a by binding to their common receptor. IL1ra has been proposed to be one of the mediators of the therapeutic effect of MSC by antagonizing IL1 effects and blocking inflammation. Luz-Crawford et al. (33) showed that IL1ra secreted by MSC acts on macrophages by inducing a polarisation toward the anti-inflammatory phenotype.

Third, we showed an impact of fibrin on the immunomodulatory properties of BMC. The secretomes from both BMC and F-BMC alleviated the gene expression of macrophages and specifically induced a polarisation of M_(LPS/INF)_ and M(_-_)toward alternatively activated macrophages phenotype. Indeed, BMC and F-BMC educated macrophages increased their expression of anti-inflammatory markers (*Arg1*, *Cd163*, *Cd206*, *Il10*, *Tgfb*) while decreasing pro-inflammatory markers (*Nos2*, *Cd86*, *Il1b*, *Il6*, *Mcp1*, *Tnfa*).

It is well established that MSCs promote the polarisation of macrophages to an anti-inflammatory phenotype (34–37). For instance, co-cultures of MSC and macrophages have shown that MSC induces the conversion of classically activated, pro-inflammatory to alternatively activated macrophages (38). Interestingly, our study indicate that the combination of fibrin and BMC further potentiated the anti-inflammatory regulatory capacity of BMC.

Alternatively-activated macrophages mediate the resolution of inflammation by phagocytosis of cellular debris, production of ECM proteins, and secretion of cytokines such as IL10 and TGF-beta (39). Modulating macrophages phenotype for salvaging ischemic damage is a promising new therapeutic strategy. Indeed, dampening the inflammatory response using MSC is broadly studied for multiple chronic diseases (40). As far as MI is concerned, several cell-based treatments increased CD206^+^ macrophages. It has been demonstrated that stimulating the anti-inflammatory CD206^+^ macrophage polarisation with cytokines, bioactive drugs or cell treatments improved cardiac tissue repair in MI animal models (41, 42). The current assumption is that a transient CD206^+^ phenotype effectively clears inflammation and may be advantageous for improved cardiac function and alleviated adverse ventricular remodelling (41). Nevertheless, a chronic elevation of CD206^+^ cells could have unwanted consequences due to the fibrotic properties of the alternatively activated macrophage population (43). Remarkably, the pro-and anti-fibrotic environment may influence the balance between the different macrophage populations identified in MI.

Furthermore, in M_(IL4)_, *Tnfa* expression was significantly downregulated by F-BMC, and its secretion remained low. Likewise, *Tnfa* secretion by M_(LPS, INF)_ and to a lesser extent by M_(-)_ was significantly reduced in F-BMC educated macrophages. Accordingly, an MSC-conditioned medium has been shown to inhibit the production of TNF-alpha by activated macrophages in vitro through the release of Il-1ra (44). Therefore, a possible explanation of this reduction is the increased of *Il1ra* in F-BMC.

In addition, F-BMC induced an upregulation of *Tgfb* in all the macrophages subsets. Notably, it has been shown that GREM1 increased the expression of *Tgfb* in hepatic stellate cells and tubular cells (45). Consistently, the increased GREM1 in the F-BMC secretome could potentially mediate the *Tgfb* elevation.

Fibrin was obtained from fibrinogen and thrombin. Accordingly, the secretome of F-BMC contains more fibrinogen than the one of BMC. Fibrinogen is thought to activate macrophage inflammatory pathways and might affect macrophage polarization in our model. Nevertheless, the pro-inflammatory effect of fibrinogen is inhibited by fibrin (46). Here, we showed a negligible pro-inflammatory impact of fibrinogen. For instance, the conditioned medium from fibrin, alone, downregulated the pro-inflammatory related genes and upregulated Cd163 in M_(IL4)_.

### Cardiomyoblast fate

F-BMC stimulated the spreading of cardiomyoblasts but not their proliferation. Notably, the proteomics analysis revealed increased Osteopontin (OPN/SSP1) in the secretome of F-BMC. ONP has been associated with cardiac hypertrophy (47). Therefore, the F-BMC induced cardiomyoblast hypertrophy is consistent with OPN level.

Macrophages had an impact on cardiomyoblasts proliferation in a phenotype dependent manner. Uneducated M_(IL4)_ greatly stimulated H9C2 proliferation while others macrophage subsets did not. Nevertheless, when educated by F-BMC, M_(LPS,INF)_ and M_(-)_ developed a capacity to stimulate cardiomyoblast growth compared to uneducated macrophages. This effect was also promoted by fibrin for M_(LPS/INF)_ and by BMC for M_(-)_. Taking all together, our results suggest that mitogenic properties developed by F-BMC educated M_(LPS/INF) and_ M_(-)_ could be a consequence of their polarisation toward an anti-inflammatory phenotype.

Altering cardiomyocyte fate emerged as an effective strategy to compensate for the loss of functional cardiomyocytes following MI. In the present study, we validate *in vitro* that the use of fibrin and BMC are potential catalyzers of optimal microenvironment conditions to favour the polarisation of macrophages. Their role in cardiac repair remains to be investigated. Psarras and al (7) identified up to seven new cardiac myeloid cell subtypes, including four macrophage populations; their respective role is still not fully understood (6). However, the cross-talk between cardiac cells and macrophages is essential for cardiac homeostasis (7, 11, 42, 48–50). In addition to their multifaced roles in ECM modulation, cardiac electrical conduction and mitochondrial homeostasis (51–53), we reinforce the importance of anti-inflammatory macrophages in regulating proliferative cell fate. The present *in vitro* results suggest that macrophage plasticity and anti-inflammatory environment are both necessary for modifying cardiomyoblast cell size and proliferation. Nevertheless, further studies need to be undertaken to investigate these effects on other models such as neonatal and adult cardiomyocytes *in vitro* and also *in vivo* and validate this present proof of concept. Indeed, Vagnozzi et al. (11) show *in vivo* no formation of new cardiomyocytes following temporal and regional induction of CCR2 ^+^ and CX3CR1^+^ macrophages. Nevertheless, the understanding of the spatiotemporal role of the macrophage in limiting adverse remodelling is still unclear. Our study suggests that (1) the importance of a scaffold that stimulates an optimal immune response for regeneration is to be considered and (2) F-BMC may potentially induces early stimulation of the cardiac reparative process. Further studies are necessary to identify the detailed mechanism *in vitro* and acute *in vivo* effects.

### Integrated concept

Taking all together, as represented in Figure 12, we documented *in vitro* that first, the F-BMC secretome has crucial effects on macrophages phenotypes: F-BMC induces the polarization of M_(-)_ toward an anti-inflammatory phenotype, a phenotype switch of M_(LPS,INF)_ and the proliferation of M_(IL4)_. Accordingly, the F-BMC educated macrophages present an anti-inflammatory phenotype and their secretomes favour cardiomyoblast proliferation. Second, the F-BMC secretome promotes cardiomyoblast spreading.

**Figure 12:**
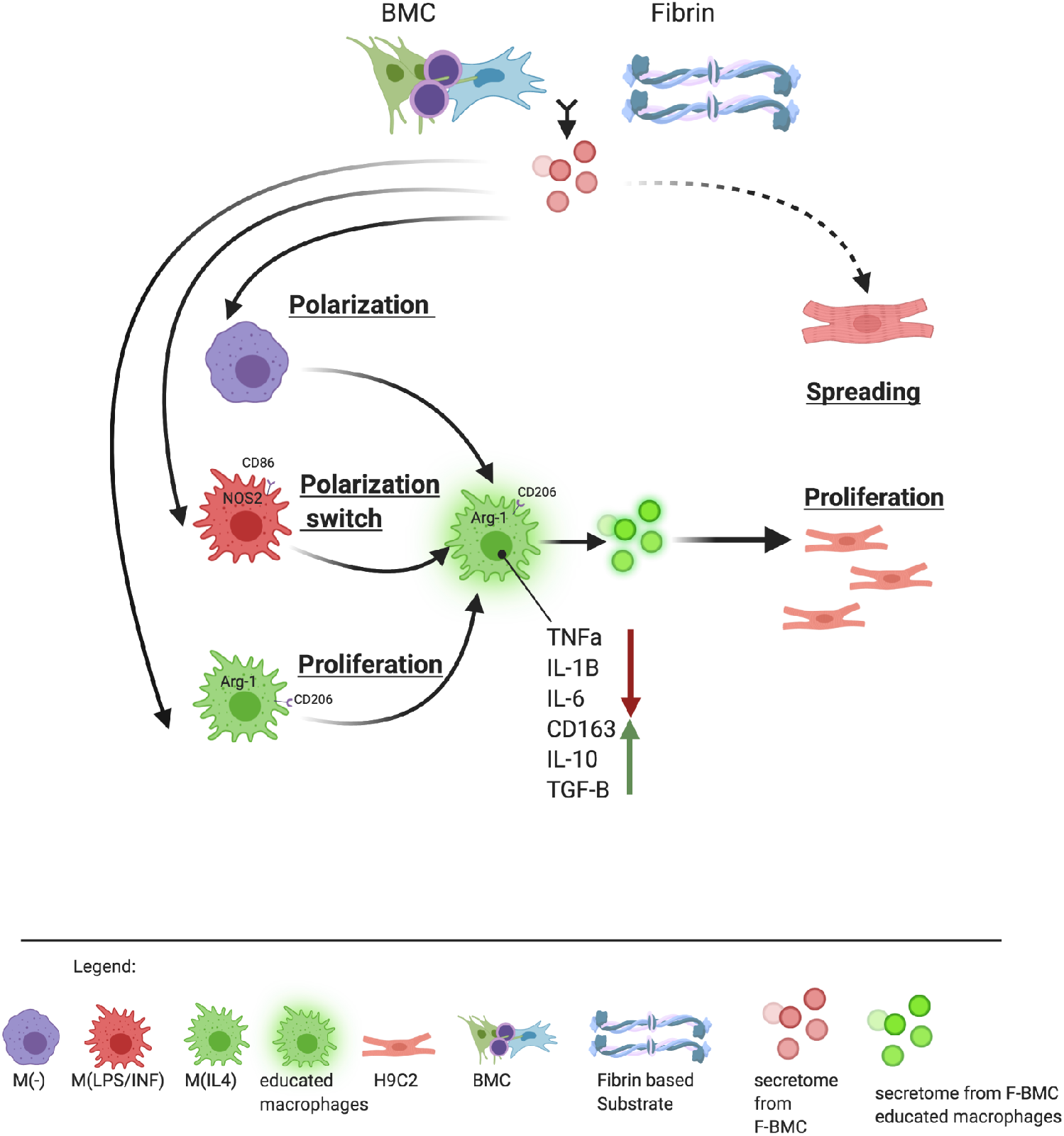
Schematic representation of the integrated concept of the effect of F-BMC on two types of cells: macrophages and cardiomyoblasts. First, the secretome of F-BMC has different effects on different macrophage subsets and induces a polarization of M_(-)_ toward an anti-inflammatory phenotype, a phenotype switch for M_(LPS,IFN)_ and stimulated M_(IL4)_ proliferation. Second, the F-BMC educated macrophages showing an anti-inflammatory phenotype increase cardiomyoblast proliferation. In parallel, F-BMC favours cardiomyoblast hypertrophy.

In conclusion, our study provides evidence that *in vivo*, F-BMC treatment lowered the infarct extent, increased wall thickness and improved cardiac function. *In vitro*, the F-BMC secretome promoted the growth of anti-inflammatory macrophages, stimulated macrophage plasticity and consequently altered the balance between the pro-and anti-inflammatory macrophage subsets. F-BMC secretome favoured the mitogenic properties of anti-inflammatory macrophages promoting cardiac cell growth.

## Methods

### In vivo study

#### Animals

Animals were purchased from Janvier (Le Genest, France) and received humane care in compliance with the European Convention on Animal Care and agreement with the Swiss Animal Protection Law. The protocol was approved by the cantonal and Swiss FederalVeterinary Office, Switzerland (FR-2016-41) and controlled for reduced animal suffering (all animals received, post-surgery, subcutaneous injection of 0.1 mg kg/1 Temgesic).

#### Bone-marrow derived cells isolation BMCs and characterization

BMCs were collected from femurs and tibias of adult rats, flushed with sterile phosphate-buffered saline (PBS; Carl Roth GmbH, Germany), incubated in red blood cell lysis buffer (Sigma, United Kingdom) and cultured in Iscove’s Modified Dulbecco’s Medium (IMDM, Pan Biotech, Germany) supplemented with 20% Fetal Bovine Serum (FBS, Biochrom AG, Berlin), 100 IU ml–1 penicillin and 100 mg ml–1 streptomycin (Corning, USA). The initial culture medium was changed at passage 1 to Dulbecco’s Modified Eagle’s Medium-high glucose (DMEM, Sigma-Aldrich, Switzerland), with 10 % FBS, 100 IU ml–1 penicillin and 100 mg ml–1 streptomycin. Cells were cultured under aseptic conditions using sterile and RNAse/DNAse free tissue culture-treated plastic ware (Corning, USA) at 37°C and 5% CO2 in a humidified incubator (Model CB160, Binder, Germany). The medium was replaced every two days, and upon 80% confluence, cells were trypsinized, pooled, and subcultured until passage 2. In order to preserve the different populations of the BMCs, cells were collected at passage two, and no other selection was performed.

For in vivo study, each pool was obtained from 10 male Lewis (mean weight of 150g). For the in vitro study, each pool of BMCs was prepared from 3 male Lewis rats (mean weight of 150g).

#### Myocardial infarction model

A total of 51 Female Lewis rats (mean weight of 200-220g) were included in the study: following a permanent left anterior descending (LAD) coronary ligation. All surgical interventions were performed under isoflurane and oxygen (5% for induction and 2.5% for maintenance), animals were placed on a warming pad at 37°C to avoid hypothermia during anaesthesia and ventilated with a 14-G IV cannula at 70-90 cycles per minute (Harvard Inspira Apparatus, Inc.; Holliston, MA, USA). Anaesthetized and ventilated as described. For surgical induction of MI, the proximal left anterior descending (LAD) coronary artery ligation was accessed through a left thoracotomy between the fourth and fifth intercostal space and was permanently ligated (7/0 polypropylene suture, Ethicon, Inc.; Somerville, MA, USA) (54). Four animals died during or a few days post MI induction. Blood samples were collected 24h post-MI from the caudal tail artery in the anaesthetized rats, and plasma was isolated using BD Vacutainer® Cell Preparation Tube (Becton Dickinson) with Sodium Heparin following manufacturer’s instructions. The plasma fraction was immediately frozen and stored at −80°C. After thawing the samples, the plasma cardiac Troponin T (cTnT) level was analyzed on a System Roche Hitachi Cobas (Roche Diagnostics, Switzerland) (55). The criterium for selecting the animals was a cTnT level > 900 ng/L (55), resulting in the exclusion of four animals. The mean plasmatic cTNT level was 3028 ± 956 ng/L. Two weeks after MI induction, cardiac function was measured by high-resolution echocardiography; reduced cardiac function assessed after two weeks significantly decreased from 70±2% to 47±8%.

#### Epicardial treatment

Two weeks post LAD ligation, 43 rats underwent a second left thoracotomy between the fifth and sixth intercostal space under general anaesthesia (54). Animals were allocated to the treated or untreated group and received either a sham operation (untreated, n=23) or epicardial implantation on the infarcted myocardium of an association of fibrin (Tisseel®, Baxter Healthcare Pty Ltd, USA) and 2 million BMC (n=20). One animal from the latest group died 11 weeks post-treatment.

#### Histological analysis

The heart was harvested, and cross-sections of 2 mm thickness from the base to the apex were obtained for systematic sampling. Each section was embedded in paraffin using a standard histological procedure. The paraffin blocks were sectioned at 5-µm intervals with a manual microtome Shandon Finesse 325 (Thermo Fischer, USA) and stained with Masson-Goldner trichrome staining. The slices were successively incubated in Mayer’s Hematoxylin (Merck AG; Zug, Switzerland), Acid Fuchsin-Ponceau (Sigma-Aldrich; Buchs, Switzerland), Phosphomolybdic Acid Orange G (Merck AG; Zug Switzerland and Sigma-Aldrich; Buchs, Switzerland), and Lichtgrün (Sigma-Aldrich; Buchs, Switzerland) solutions. The samples were dehydrated with an ascending ethanol series and mounted with Eukitt (EM Sciences; Hatfield, PA, USA). Images were acquired with a stereomicroscope Nikon SMXZ800 mounted with a Nikon 1 camera (Nikon; Tokyo, Japan). Bersoft Image Analysis software (Bersoft Technology and Software; Lunenburg, Canada) was used to measure the scar thickness of the infarct, septum thickness, left ventricle (LV) cavity area, infarct area, and LV tissue area. The infarct expansion index (EI) was calculated as ((LV cavity area/whole LV area)/ (infarct thickness/septum thickness)). The measurements were performed on one 5-µm slice from each 2 mm heart section. EI for each heart was the average of 5-6 sections (56).

### In vitro studies

#### Conditioned medium preparation

BMCs (0.3×10^6^ cells) were cultured in 6 well plates coated with 40 uL fibrin (20 ul Thrombin and 20 ul fibrinogen; Tisseel®, Baxter Healthcare Pty Ltd, USA) (F-BMC) or in an uncoated plate (BMC). The ratio cell/fibrin was optimized for low cell mortality (<3% when measured with Propidium Iodine). Fibrin conditioned medium was obtained after 48h from a plate coated with 40 uL fibrin. Cd-medium were collected and centrifuged for 5 min at 1200 rpm, immediately used or stored at -80°C for proteomic analysis.

#### MS-based proteomics

Cd-medium was concentrated by ultrafiltration using vivaspin columns (10 kDa MWCO). Samples were heated in SDS-PAGE loading buffer, reduced with 1 mM DTT for 10 min at 75°C and alkylated using 5.5 mM iodoacetamide for 10 min at RT. Protein mixtures were separated on 4-12% gradient gels (Nupage, Thermo Fisher). Gel lanes were cut into 6 slices, and proteins therein were in-gel digested with trypsin (Promega, Dübendorf, Switzerland), and resulting peptide mixtures were processed on STAGE tips and analyzed by LC-MS/MS.

Mass spectrometric measurements were performed on a QExactive Plus mass spectrometer (Thermo Scientific) coupled to an EasyLC 1000 nanoflow–HPLC. HPLC–column tips (fused silica) with 75 µm inner diameter were self-packed with Reprosil–Pur 120 C18-AQ, 1.9 µm (Dr Maisch GmbH, Ammerbuch, Germany) to a length of 20 cm. Samples were applied directly onto the column without a pre-column. A gradient of A (0.1% formic acid in water) and B (0.1% formic acid in 80% acetonitrile in water) with increasing organic proportion was used for peptide separation (loading of the sample with 0% B; separation ramp: from 5–30% B within 85 min). The flow rate was 250 nL/min and for sample application 600 nL/min. The mass spectrometer was operated in the data-dependent mode and switched automatically between MS (max. of 1×10^6^ ions) and MS/MS. Each MS scan was followed by a maximum of ten MS/MS scans using a normalized collision energy of 25% and a target value of 1000. Parent ions with a charge state from z = 1 and unassigned charge states were excluded for fragmentation. The mass range for MS was m/z = 370–1750. The resolution for MS was set to 70,000 and for MS/MS for 17,500. MS parameters were as follows: spray voltage 2.3 kV; no sheath and auxiliary gas flow; ion–transfer tube temperature 250°C.

MS raw data files were uploaded into the MaxQuant software version 1.6.10.43 for peak detection, generation of peak lists of mass error-corrected peptides, and for database searches (PMID 19029910). A full-length UniProt rat database additionally containing common contaminants such as keratins and enzymes used for in-gel digestion (based on UniProt rat FASTA version May 2019) was used as reference. Carbamidomethylcysteine was set as fixed modification, and protein amino-terminal acetylation and oxidation of methionine were set as variable modifications. LFQ was chosen as the quantitation mode. Three missed cleavages were allowed, enzyme specificity was trypsin/P, and the MS/MS tolerance was set to 20 ppm. The average mass precision of identified peptides was, in general, less than 1 ppm after recalibration. Peptide lists were further used by MaxQuant to identify and relatively quantify proteins using the following parameters: peptide and protein false discovery rates, based on a forward-reverse database, were set to 0.01, minimum peptide length was set to 7, the minimum number of peptides for identification and quantitation of proteins was set to one which must be unique, minimum ratio count was set to two, and identified proteins were again quantified. The interleukins were not in the detection range of the actual proteomic protocol.

#### Macrophage isolation, differentiation and priming with condition media

Macrophages were isolated from the bone marrow of seven different pools of 3 male Lewis rats (mean weight of 150g) (Janvier, France) and cultured during seven days in DMEM medium, 10 % FBS, 100 IU ml–1 penicillin and 100 mg ml–1 streptomycin, supplemented with 50 ng ml^−1^ of macrophage colony-stimulating factor (M-CSF, Peprotech, London, United Kingdom) (Murray 2014). Cells were washed with PBS and stimulated with either LPS (50 ng ml^−1^; Sigma-Aldrich; Buchs, Switzerland) and IFN (10 ng ml^−1^; rat recombinant; Peprotech, London, United Kingdom) to trigger their differentiation towards a pro-inflammatory phenotype M_(LPS,IFN)_ or IL4 (20 ng ml^−1^, rat recombinant; Peprotech, London, United Kingdom) to obtain anti-inflammatory macrophages, M_(IL4)_ or kept untreated for unpolarised macrophages (M_(-)_)(adapted from (38)). M_(-)_, M_(LPS,INF)_ and M_(IL4)_ were further cultured, during 48 hours, with F-BMC, BMC or fibrin conditioned media. Culture with DMEM 10% medium served as control. The differentiation of the macrophages was validated by surface markers and expression profiles accessed by immunostaining and RT-PCR, respectively.

#### Enzyme-linked immunosorbent assay (ELISA)

The macrophages were cultured with F-BMC, BMC or substrate conditioned media, washed with PBS and then cultured with standard medium for 48h. All cytokines were quantitated from cell culture supernatants of the different macrophages phenotypes. ELISA kits (IL-1β (ab100767; Abcam, United Kingdom), IL-6 (ab119548; Abcam, United Kingdom), TNF-alfa (ab46070; Abcam, United Kingdom) were used according to the manufacturer’s instructions and were estimated as pg/mL. Absorbance was read on Tecan Infinite 200 PRO Microplate Reader (Tecan Group, Switzerland).

#### Real-time Polymerase chain reaction

RNA was isolated from BMC cultured with or without fibrin and from the different macrophages phenotypes previously cultured with BMC, F-BMC, fibrin conditioned media or with standard growth medium, using the Trizol Reagent (Molecular Research Center, Inc., Cincinnati, OH, USA) according to the manufacturer’s instructions. RNA was reverse transcribed to generate complementary DNA, using GoScript Reverse Transcription Mix™ (Promega, USA) according to the manufacturer. Two-step quantitative Real Time-PCR was performed to measure mRNA expression using StepOne SYBR System (Thermo Fisher Scientific, Switzerland) with GoTaq® qPCR Master Mix (Promega, USA) and acquired with Step One software v.2.3. (Thermo Fisher Scientific, Switzerland). mRNA expression was assessed using primers presented in Supplementary Table S2. The mRNA expression levels of all genes were quantified by normalizing to the geometric mean of the reference genes *Gapdh* and *beta-Actin* and using the relative quantification of gene expression calculated by the 2^−ΔΔCt^ approximation method.

#### H9C2 rat cardiomyoblasts

H9C2 rat cardiomyoblasts (Sigma-Aldrich; Buchs, Switzerland) were expanded in Dulbecco’s Modified Eagle’s Medium-high glucose (DMEM, Sigma-Aldrich, Switzerland), with 10 % FBS, 100 IU ml–1 penicillin and 100 mg ml–1 streptomycin. Cardiomyoblasts were further cultured during 48h with DMEM 10% medium or conditioned media from F-BMC, BMC, fibrin, or macrophages. Additionally, H9C2 were cultured with F-BMC, fibrin, or BMC educated macrophages. To obtain the double conditioned medium, the macrophages were cultured with the fibrin, BMC, or F-BMC conditioned media for 48h; the educated macrophages condition media were collected, centrifuged and added to the H9C2.

#### Real-time cell analyzer system (RTCA) and EdU cell proliferation assays

10000 cells/well were seeded in an E-Plate® with gold microelectrodes fused to the bottom surface and cultured for 120 hours using the instrument xCELLigence Real-Time Cell Analyzer (RTCA; ACEA Biosciences Inc, Germany) according to the manufacturer’s instructions. The cell index (CI; impedance measurement correlated with cell proliferation) was recorded for 120 hours. CI was plotted against time (hours), and the area under the curve (AUC) was calculated. (Moniri et al., 2015).

EdU Cell Proliferation Kit (Sigma-Aldrich; Buchs, Switzerland) was used according to the manufacturer’s instructions. Pictures of stained cells were acquired using Cytation 5 Cell imaging Multi-mode reader (Biotek, Switzerland), and the percentage of positive EdU cells per field was counted using Gen5 software (Biotek, Switzerland).

#### Statistical analysis

Statistical analyses were performed using statistical software GraphPad Prism, Version 8 (GraphPad Software, San Diego, CA, USA). All values were reported as mean ± standard deviation (SD). Data distribution was assessed using the Kolmogorov-Smirnov test. For the in vivo study, an unpaired t-test was performed. For the in vitro study, one-way ANOVA was performed, followed by a Dunnett’s multiple comparisons test or otherwise indicated. Results were considered significant from *p* < 0.05.

## Acknowledgements

The study was supported by the University of Fribourg, the Swiss National Science Foundation (SNF 310030_149986) attributed to MNG, and the Fonds Scientifique Cardiovasculaire FSC, Fribourg Hospital attributed to SC. The proteomic analysis was part of the SKINTEGRITY.CH collaborative research project supported by the University of Fribourg (JD).

## Author contributions statement

IB and MNG designed the experiments; IB, JV, AF, GA and BF conducted the experiments; MNG, IB and JD analyzed the results; IB and MNG prepared the figures; IB and MNG wrote the main manuscript. The authors reviewed the manuscript.

## Data availability

The datasets generated and analyzed during the current study are available from the corresponding author on reasonable request.

## Supplementary data

**Figure S1:**
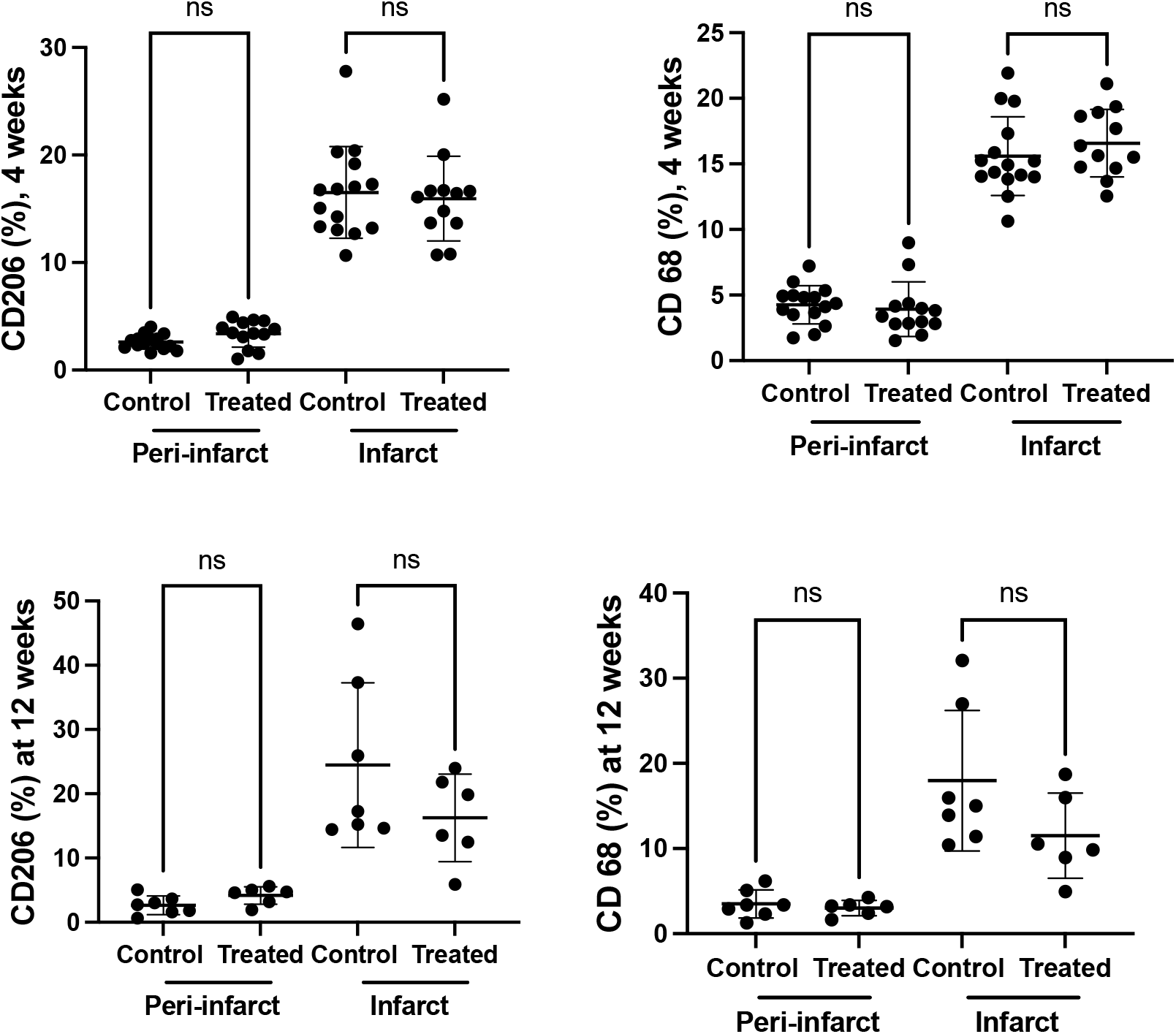
CD68+, CD206+ macrophages within the infarcted area and border zone were similar in all groups. Immunostainings for CD206 (A, C) and CD68 (B, D) were performed on paraffin-embedded heart sections. 5 to 6 cross-sections from a systematic sampling of the whole heart were averaged for each animal. Hearts were harvested 4 or 12 weeks post-treatment.

### Methods

Macrophages immunohistochemistry: Samples from rat hearts sectioned into 5-6 parts starting at the base and going to the apex were used for double fluorescent immunochemistry staining. Anti-CD206 (Abcam, ab64693) and anti-CD68 (Abcam, ab31630) were used as primary antibodies and Donkey anti-Rabbit Alexa Fluor® 647 (Abcam, ab150075) and Goat anti-mouse Alexa Fluor® 488 (Abcam, ab150113) were used as secondary antibodies. The tissue sections were counterstained with Hoechst. Images of the left ventricle were acquired using a DM6B bright-field microscope (Leica) at 10x magnification. The quantification was done on FIJJ (ImageJ). Two regions within the infarct zone and the adjacent infarct zone were delimited for each image. An auto-threshold was performed. On each region of interest (ROI), the number of CD206 and CD68 positive cells, Hoechst stained nuclei and the total area covered by these cells were automatically counted. Each image’s masks were inspected manually to verify the automatic counting. CD206^+^ and CD68^+^ cells ratio was calculated by dividing the number of CD206^+^ or CD68^+^ cells by the total nuclei for each ROI.

---

**Figure S2:**
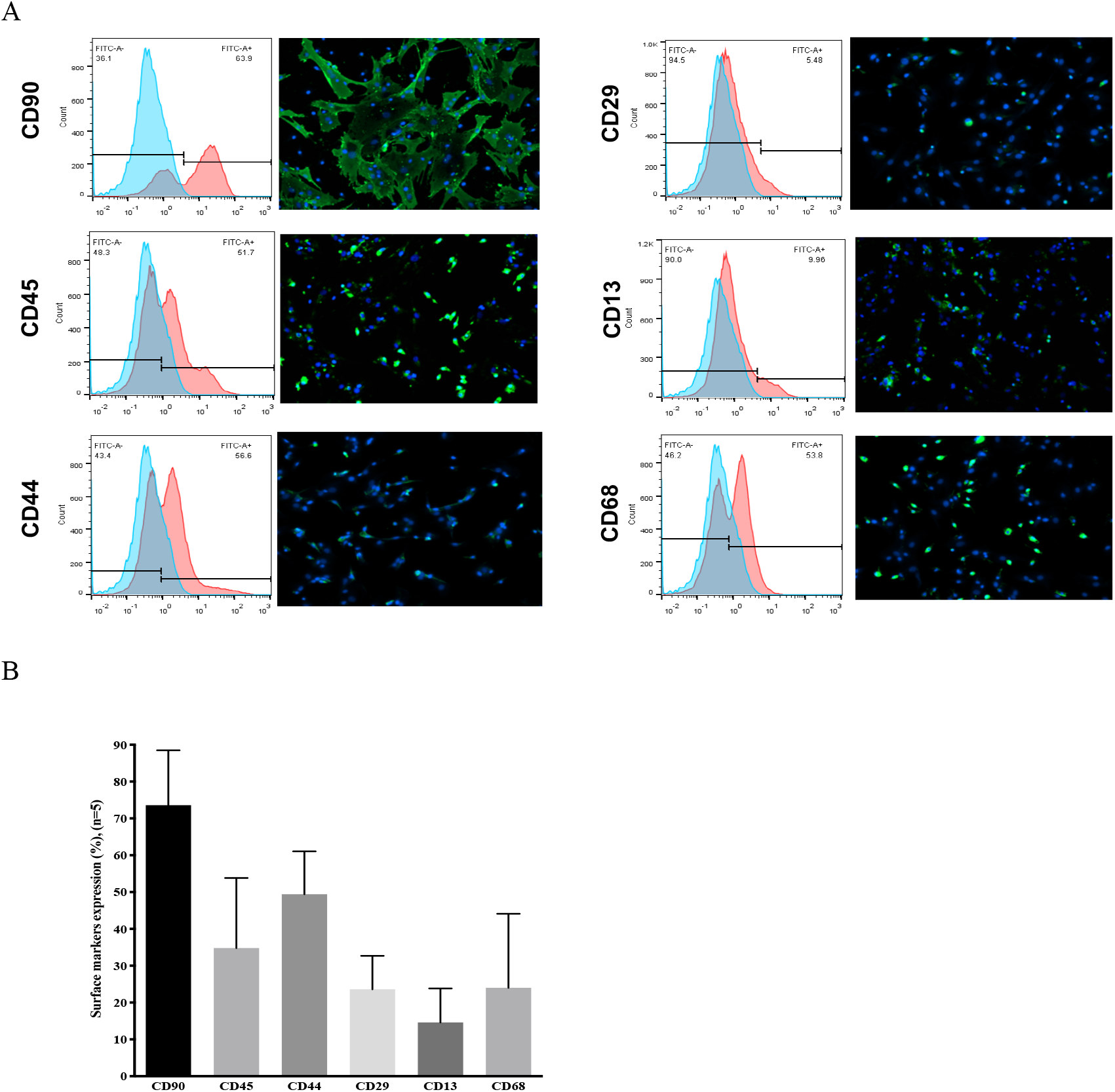
Surface markers of BMCs measured by flow cytometry and immunostaining. (A) Histogram represents the flow cytometry results, and the pictures represent the immunostainings of the cells. BMC are a heterogeneous population of cells as indicated by markers from the mesenchymal (CD90, CD29, CD13) and hematopoietic (CD45, CD44, CD68) lineages. The mesenchymal CD90^+^ cells were most abundant. They form a population of large and spread cells. (B) Quantification of the immunostainings

## Methods

Five different pools of BMC (pools of 10 or 3 animals) used in this study were characterised by flow cytometry. Passage 2 cells were detached with accutase. Cell surface markers were analysed using unlabelled Anti-CD90 (rabbit; ab225; Abcam, United Kingdom), anti-CD45 (ab10558, RRID:AB_442810, Abcam, United Kingdom), anti-CD44 (ab119348, RRID:AB_10902529, Abcam, United Kingdom), anti-CD29 (ab179471;

RRID:AB_2773020, Abcam, United Kingdom), anti-CD13 (ab108310, RRID:AB_10866195, Abcam, United Kingdom) and anti-CD68 (ab31630, RRID:AB_1141557, Abcam, United Kingdom) in 1:50 dilution. Cells were further labelled with a secondary antibody (1:500 dilution), including Alexa Fluor 488-conjugated anti-rabbit (ab150077, RRID: AB_2630356, Abcam, United Kingdom), Chromeo 488-conjugated anti-mouse (ab60313, RRID: AB_954967, Abcam, United Kingdom) and FITC-conjugated anti-rat (ab6730, RRID: AB_955327, Abcam, United Kingdom). Stained cells were fixed with 1% Para-Formaldehyde and stored at 4 °C until further analysis. Negative controls without primary antibodies were included. Flow cytometry was performed, and cell surface markers expression was acquired on a MACSQuant® Analyser 10 (Miltenyl Biotec, Germany) instrument, and the data were analysed using the FlowJo 10.4.2 (Flowjo Cloud, USA) software.

Furthermore, BMC cultured in tissue culture chamber (Sarsted, Mümbrecht, Germany) and fixed with paraformaldehyde 4% were characterised by immunostaining for CD90, CD45, CD44, CD29, CD13, CD68 according to the procedure described for flow cytometry. Cells were counterstained with Hoechst (1:200 dilution; Sigma-Aldrich, Switzerland) and examined under a high-resolution bright-field light microscope Nikon Eclipse Ni (Nikon, Japan), captured using a Nikon Digital Sight DS – Ri 1 camera and analysed by Nikon NIS elements software (Nikon, Japan).

**Figure S3:**
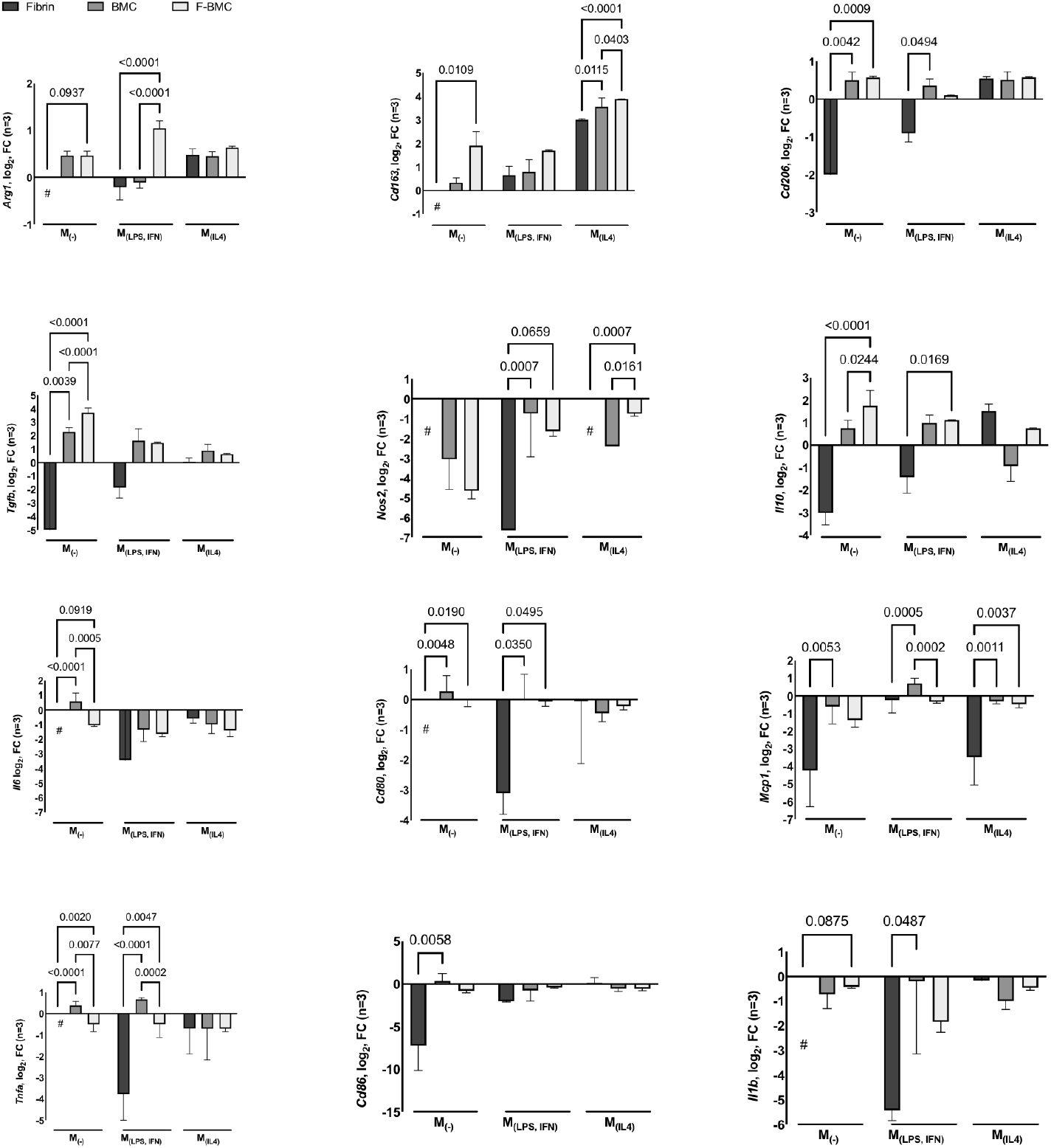
Comparison of the expression profiles of macrophages cultured in F-BMC, BMC or fibrin conditioned medium. Fold changes in gene expressions of pro-and anti-inflammatory markers were calculated for educated macrophages (M_(-)_, M_(LPS, IFN)_ or M_(Il4)_) relative to uneducated ones. The values shown are mean ± SD. All n= 3 biologically independent samples constituted of macrophages pools, each pool was obtained from 3 animals, therefore, 9 animals per group. Statistics were performed using two-way ANOVA and a turkey’s multiple comparison test as a post hoc test to compare a single effect for each macrophage subset and each gene. For M_(IL4)_, the downregulation of pro-inflammatory and upregulation of anti-inflammatory genes were similar when educated with F-BMC, BMC or fibrin except for *Cd163*, *Nos2 and Mcp1*. For M_(-)_ and M_(LPS, IFN),_ F-BMC induced significantly pronounced changes of *Arg1*, *Il10*, Cd80, *Tnfa* compared to fibrin. When compared to BMC, the significative changes are macrophage subset dependent. Significant differences were measured in M_(-)_ for *TGFb*, *Il10*, *IL6* and *TNFa.* For M_(LPS, IFN), the_ most pronounced effects were observed for *Arg1* and *Tnfa.* Notably, fibrin induced downregulation of pro-and anti-inflammatory genes principally.

**Supplemental Table 1:**
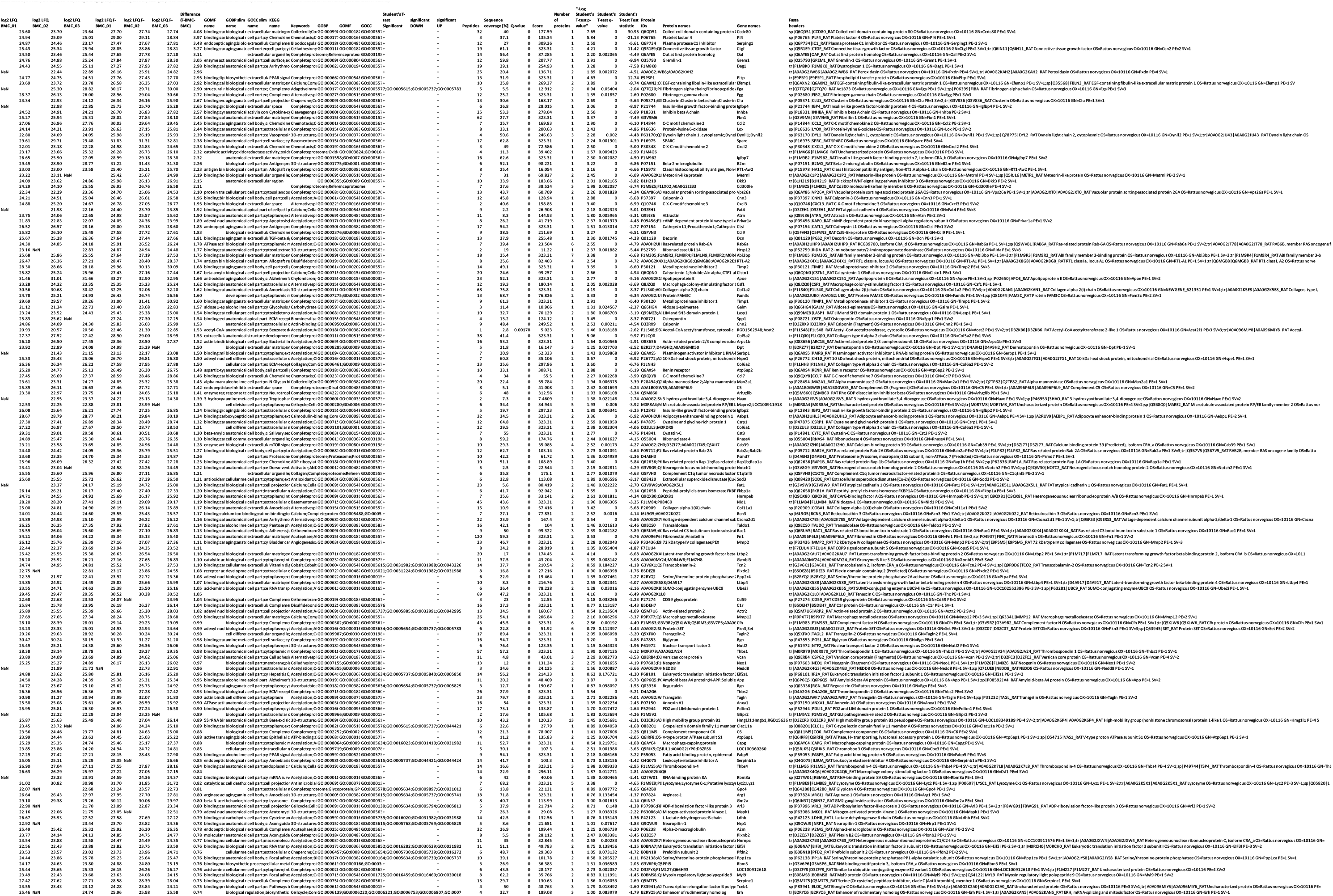
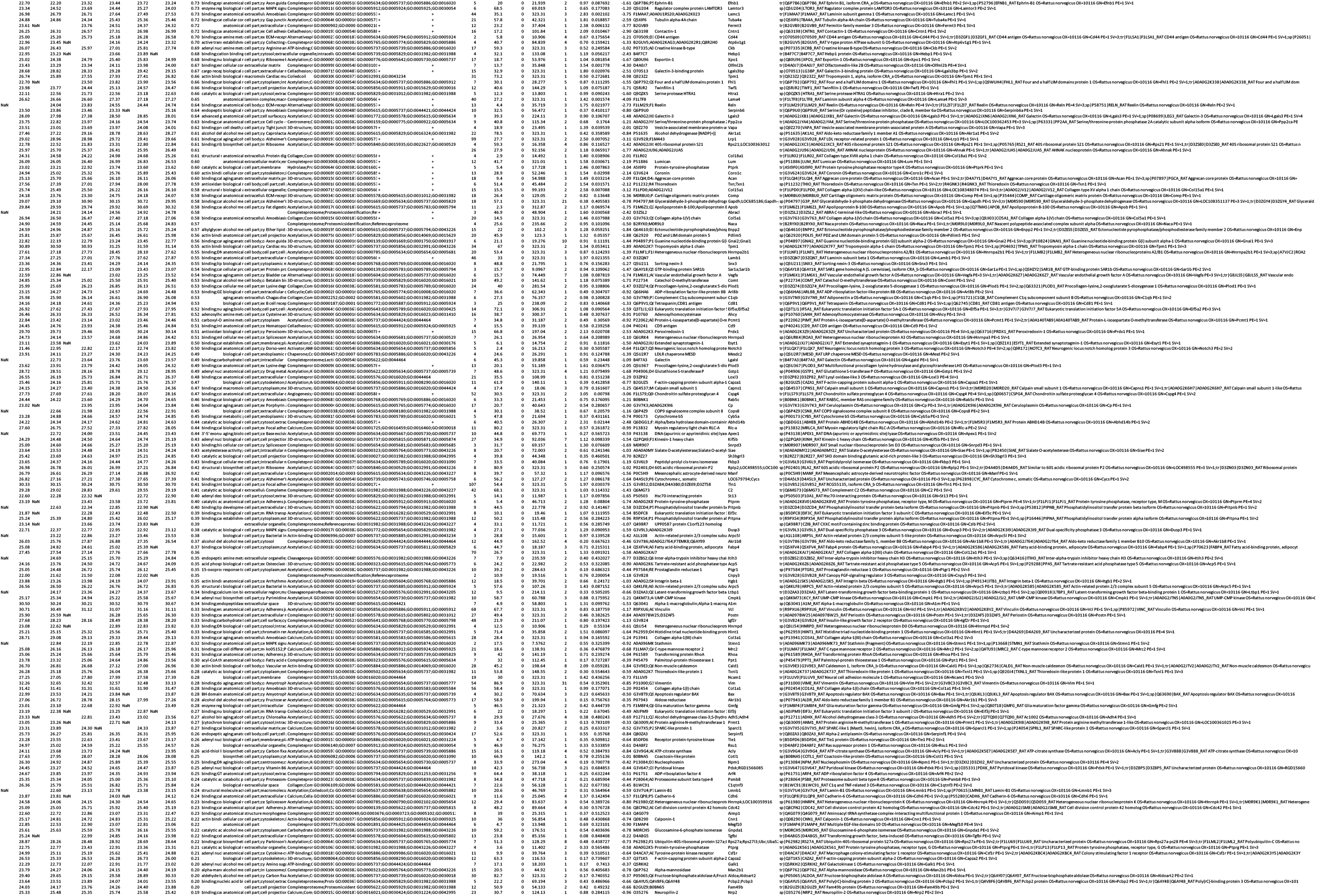
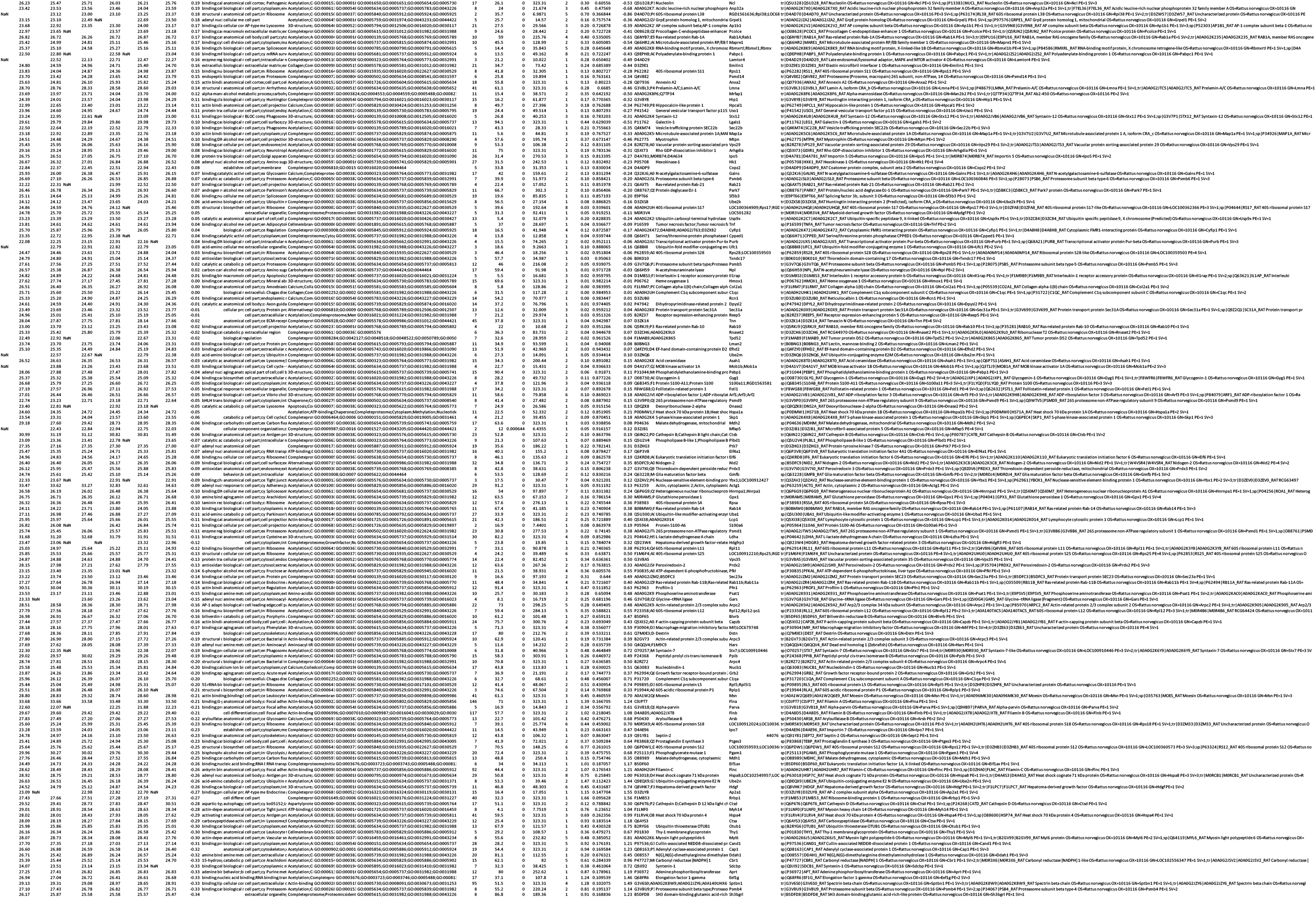
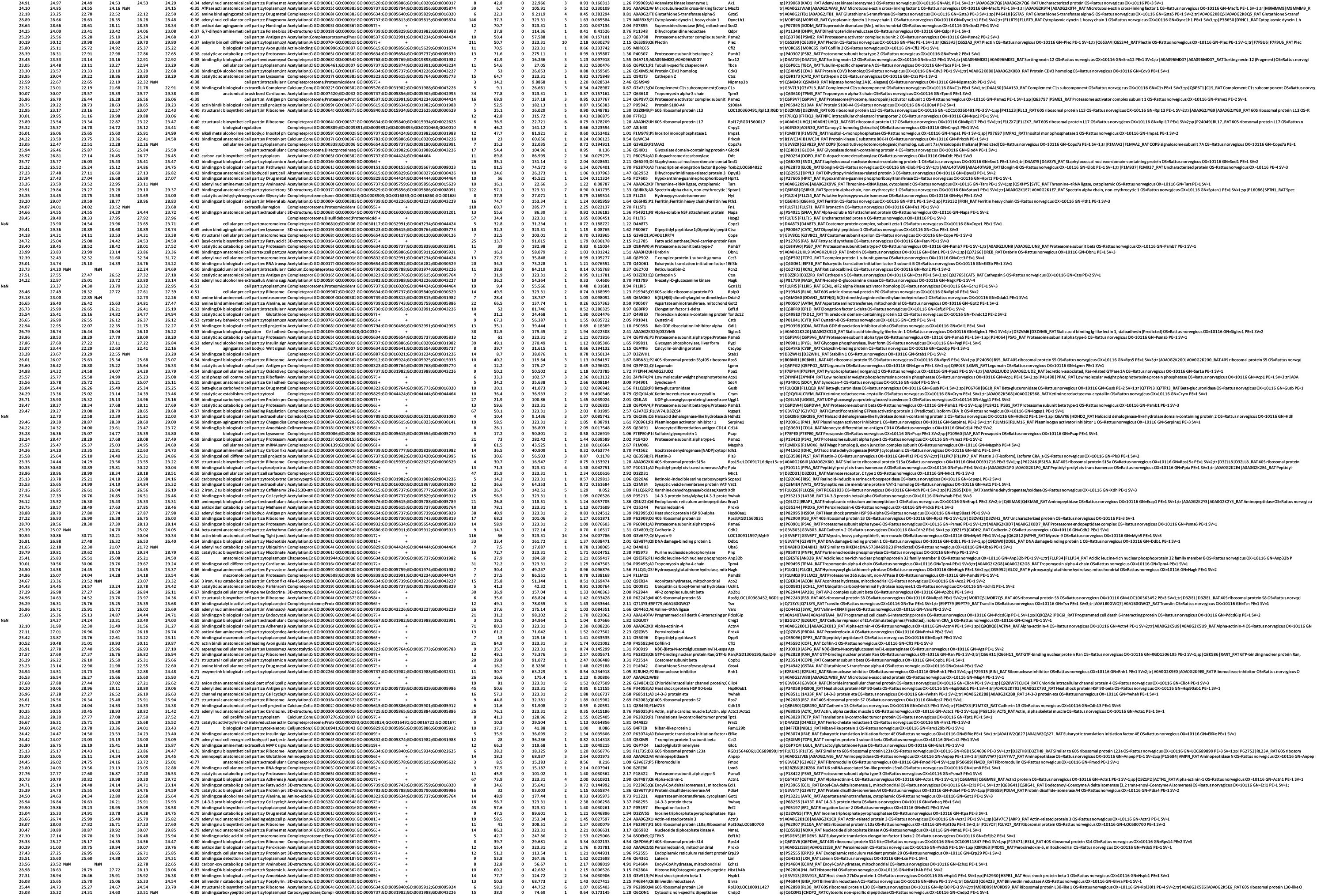
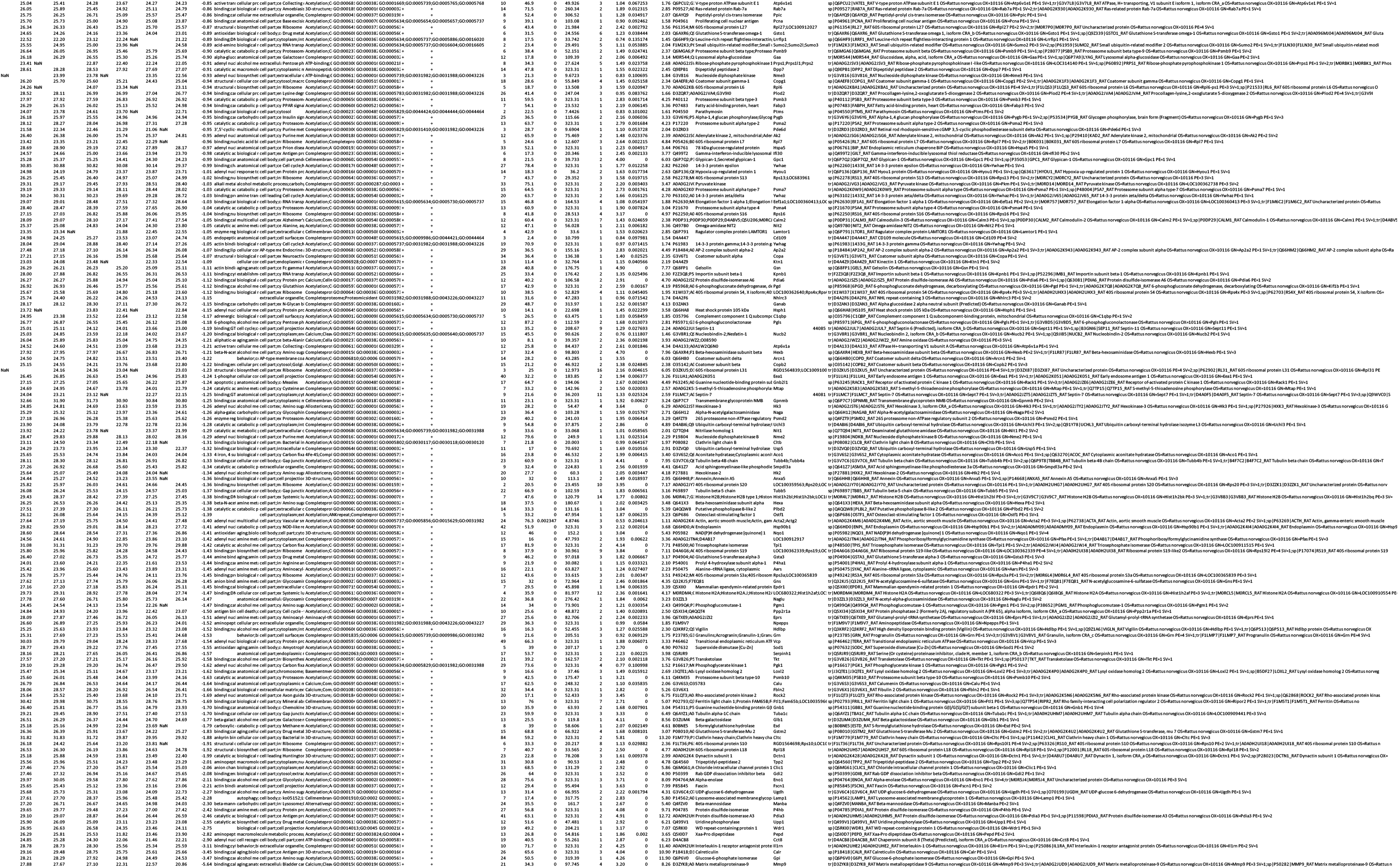
Expression proteomic data of conditioned medium of F-BMC and BMC. Three biological replicates each were analyzed by standard bottom-up proteomics. Label-free quantification (LFQ) data were log2 transformed and filtered on minimally 2 valid values per experimental group. Significant differences between both groups were calculated by a t test (FDR<0.05, S0=0.1) using average group values. Signifcantly changed proteins are marked by a “+” in columns P-R. Peptides and protein IDs were filtered on a FDR<0.01 (see Methods for details).

**Table S2:**
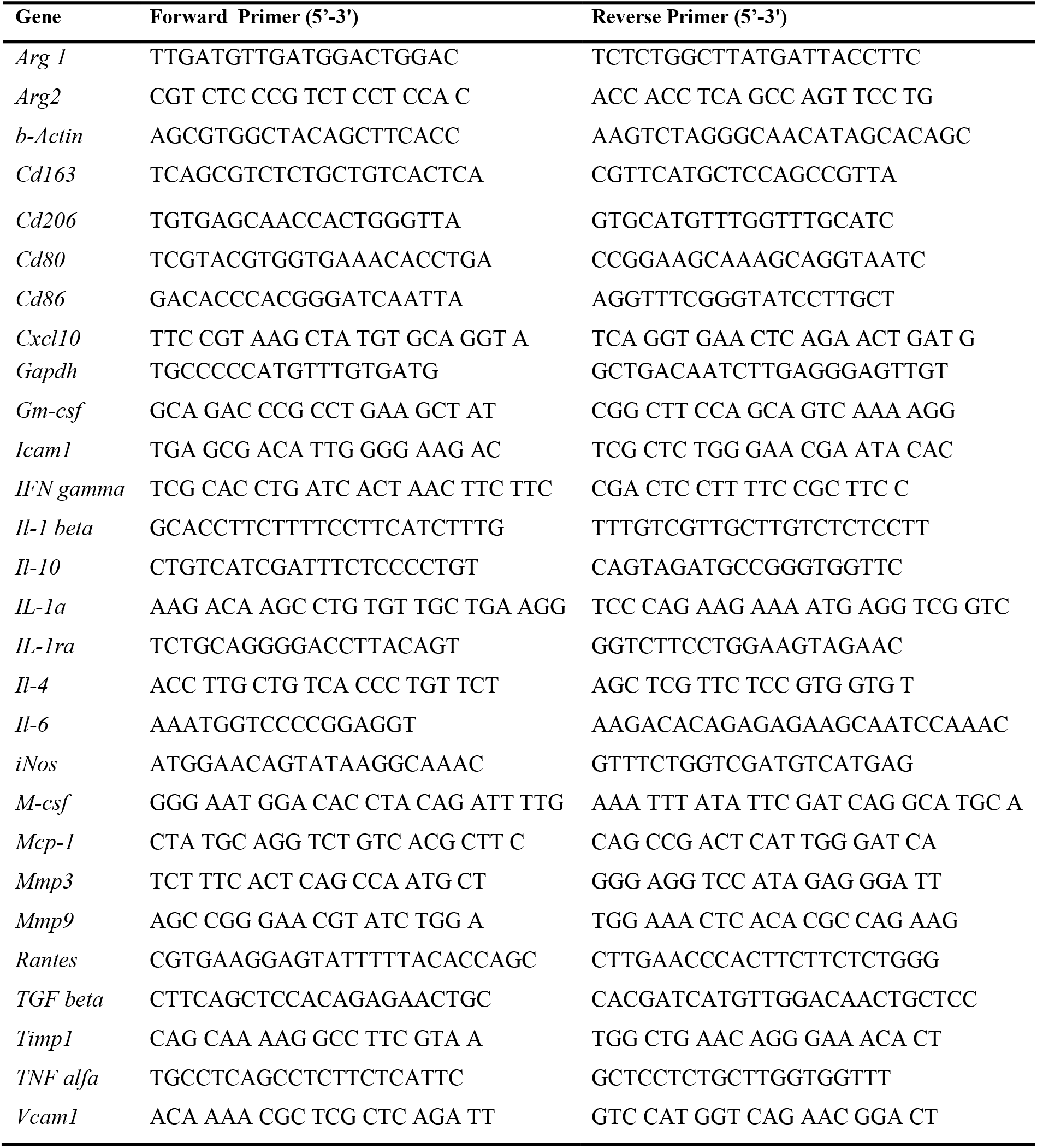
Primers used for real-time PCR analysis

